# Role of X chromosome and dosage compensation mechanisms in complex trait genetics

**DOI:** 10.1101/2025.01.16.633321

**Authors:** Yu Fu, Aino Kenttämies, Sanni Ruotsalainen, Matti Pirinen, Taru Tukiainen

## Abstract

The X chromosome (chrX) is often excluded from genome-wide association studies due to its unique biology complicating the analysis and interpretation of genetic data. Consequently, the influence of chrX to human complex traits remains debated. Here, we systematically assessed the relevance of chrX and the effect of its biology on complex traits by analyzing 48 quantitative traits in 343,695 individuals in UK Biobank with replication in 412,181 individuals from FinnGen. We show that, in the general population, chrX contributes to complex trait heritability at a rate of 3% of the autosomal heritability, consistent with the amount of genetic variation observed in chrX. We find that a pronounced male-bias in chrX heritability supports the presence of near-complete dosage compensation between sexes through X chromosome inactivation (XCI). However, we also find subtle yet plausible evidence of escape from XCI contributing to human height. Assuming full XCI, the observed chrX contribution to complex trait heritability in both sexes is greater than expected given presence of only a single active copy of chrX, mirroring potential dosage compensation between chrX and the autosomes. We find this enhanced contribution attributable to systematically larger active allele effects from chrX compared to autosomes in both sexes, independent of allele frequency and variant deleteriousness. Together these findings support a model where the two dosage compensation mechanisms work in concert to balance the influence of the chrX across the population while preserving sex-specific differences at a manageable level. Overall, our study advocates for more comprehensive locus discovery efforts in chrX.

## Introduction

Genome-wide association studies (GWAS) have discovered numerous autosomal variants associated with complex traits and diseases. However, the discovery of phenotype-associated X chromosome (chrX) loci is significantly lagging behind autosomes, despite chrX constituting ∼5% of the human genome and harboring at least 800 protein-coding genes. Indeed, it was estimated that only 25% of the published GWAS reported a chrX analysis in NHGRI-EBI GWAS Catalog^1^ as in 2021^2^, thus leaving the contribution of chrX to genetics of complex phenotypes largely unexplored. Nevertheless, genetic studies that have analyzed chrX have showcased its non-negligible role in many complex phenotypes and that novel biological discoveries can be uncovered from this chromosome^3–9^.

One of the major contributors to the exclusion of chrX has been the analytical and interpretational challenges posed by the unique biology of chrX^2,10^. Unlike autosomes that occur in pairs, in most mammals, including humans, chrX is present as two copies in genetic females (XX karyotype) but only as one copy in genetic males (XY karyotype). This leaves the extensive non-pseudoautosomal region (non-PAR) of chrX hemizygous in males. To counter the putative dosage imbalance, chrX-specific regulatory processes act to compensate for the differences in chrX dosage between males and females as well as between chrX and autosomes. Ohno proposed in 1967^11^ that dosage compensation is initiated during embryogenesis and functions through two mechanisms: (1) random X chromosome inactivation (XCI) in each female somatic cell to equalize the active dosage between sexes, leaving chrX functionally hemizygous also in XX cells, and (2) two-fold upregulation of X-linked gene expression compared to autosomal genes to balance the dosage difference between one active chrX and a pair of active autosomes.

Since first hypothesized in the 1960s by Mary Lyon^12^, XCI has now been accepted as a fact^13^ resulting in broadly equal levels of gene expression between sexes^14,15^. However, in humans, as many as 25% of chrX genes escape from XCI and continue to be expressed at attenuated level from the inactive chrX^15–17^. In contrast to XCI, the evidence supporting X-upregulation is more conflicting. Gene expression evidence generally converges to proposing partial transcriptional upregulation of chrX genes across various organisms including humans^18–23^, in a manner where the expression form a single copy of chrX is greater than that from a single autosome but lower than that from an autosome pair. Whether a similar compensatory process between chrX and autosomes extends to other biological layers, including genetic effects on complex traits, remains unclear.

Large-scale genetic data has the potential to elaborate on the male-to-female and chrX-to-autosome relationships in human complex traits in the light of dosage compensation. Full XCI is expected to manifest as a two-fold additive genetic variance in males compared to females^24^. Accordingly, comparisons of chrX SNP-heritabilities (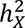) between males and females have generally supported the presence of XCI across diverse complex traits^4,5^. Escape from XCI, is, theoretically, expected to modify this relationship and lead to subtle increase in female 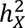 and female genetic effects. However, contrasting the transcriptome-level evidence of widespread escape^15,16,25^, genetic studies, that usually assumed complete absence of XCI at escape loci, have provided limited support for the contribution of escape to human complex traits^4^. Genetic studies that compared chrX and autosomes have found the effects of chrX to be smaller than those of a pair of autosomes in females^26^ but comparable to those of a single autosome in males^4^. Most GWAS tools nowadays support chrX analyses, facilitating the examination of chrX contributions to human complex traits. However, the inference about dosage compensation mechanisms is complicated by differing assumptions underlying these tools regarding male-to-female and chrX-to-autosome relationships (see Material and methods).

In this study, we addressed the complication in understanding chrX GWAS results due to the unique biology of chrX. We leveraged data on genotypes and 48 complex traits from 159,112 males and 184,583 females in UK Biobank (UKB)^27^ with replication data from 181,871 males and 230,310 females in FinnGen^28^ (release 10). We surveyed the contribution of chrX across the complex traits in the overall population and within each sex through partitioning SNP heritability between autosomes and chrX. We extended our study to understand how the unique biology of chrX is reflected in the phenotypic associations, through comparison of sex bias in heritabilities and genetic effects in chrX versus the autosomes. Altogether, we provide insight into the effects of chrX-specific biology on GWAS and the importance of accounting the unique features of chrX in the analysis and interpretation of genetic studies.

## Material and methods

### Complications and consequences of chrX biology in GWAS

The analysis of non-PAR in chrX poses several analytical challenges due to differing copy numbers of chrX between sexes and XCI in females. Here, we first explain the motivation and consequence of the most commonly adopted system in GWAS tools^29–32^ that codes female genotypes as {0,1,2} and male as {0,2} (other coding systems are explained in Supplemental Note) in non-PAR. Then, we discuss the power bias in sex-specific and sex-combined GWAS introduced by different copy numbers of chrX between sexes. Last, we discuss the expectations of genetic variance, heritability and effect sizes considering different degrees of escape from XCI. The effect sizes are determined through a linear regression model *Y*∼*Xβ* + *covariates*, where trait *Y* is regressed on genotype *X* to estimate the effect size *β*. We use *a* to denote the per active allele effect.

Random XCI in females results in ∼50% cells with maternal chrX active and ∼50% with paternal chrX active. Thus, assuming full XCI in females, homozygote loci (aa or AA) in a female cell are functionally equal to a hemizygous male cell with the same allele, while heterozygote loci (aA) typically have allele A functionally active in ∼50% cells and allele a in the other ∼50% of the cells.

The linear model effect sizes in males (*β*_*m*_) and females (*β*_*f*_) are equal if we assume full XCI and equal active allele effect sizes (*a*) between the sexes. When both sexes are analyzed together, we are implicitly assuming that one of the alleles in females is fully inactivated ^33^ and the estimated effect size parameter denotes half the effect of an active allele (i.e. *β*_*m*_ = *β*_*f*_ = *a*_*X*_/2). When the functionally haploid chrX causes similar magnitude of phenotypic effect as a pair of the autosome (X = AA), the estimated effects between variants in chrX and in an autosome are expected to be equal. In contrast, if one active chrX causes similar magnitude of effect as a single autosome (X = A), the estimated effects of variants in chrX are expected to be half of those of autosomal variants. We summarized the relationship between males and females, and chrX and autosomes, for different quantities in Table 1 and Figure S1.

**Table 1.**
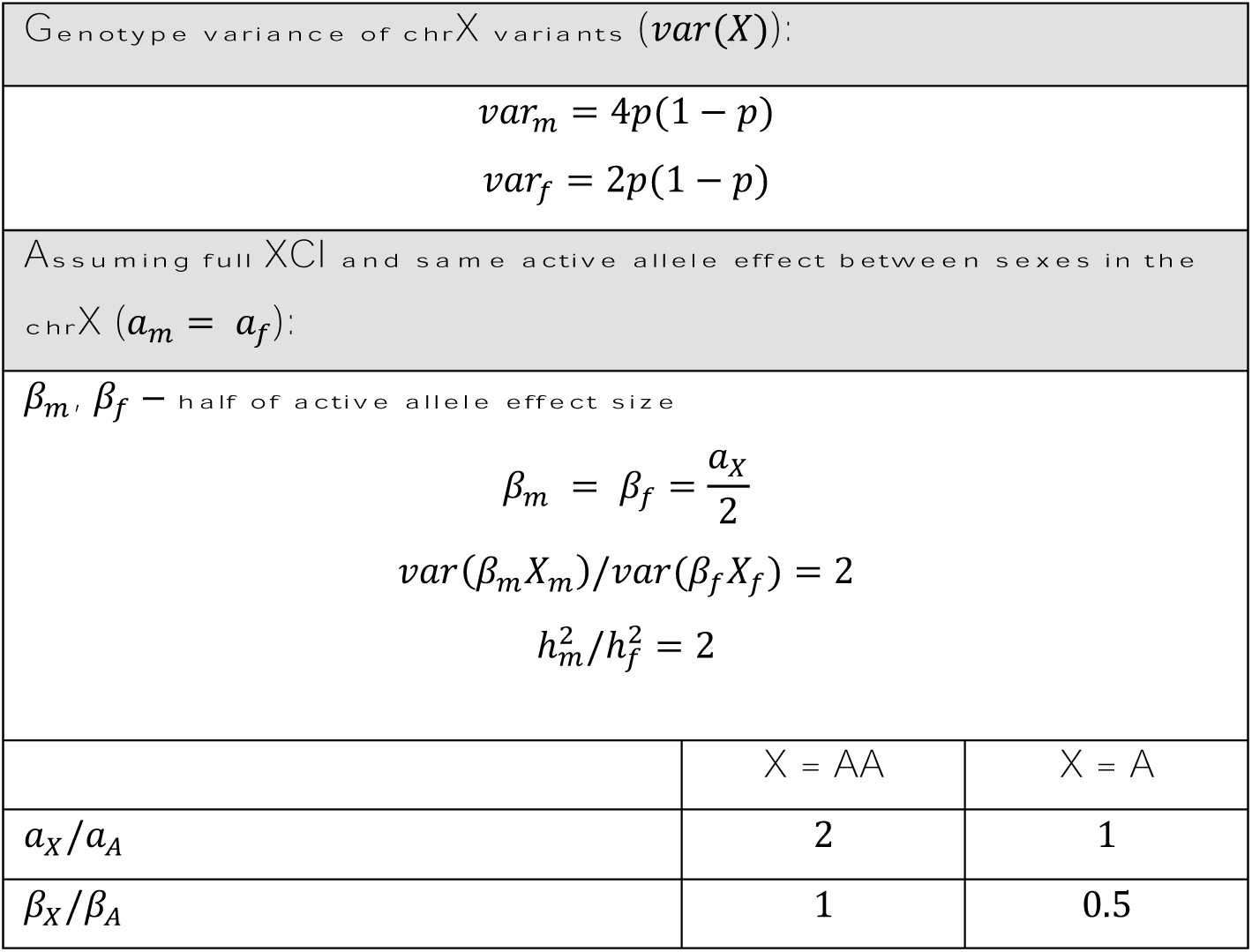
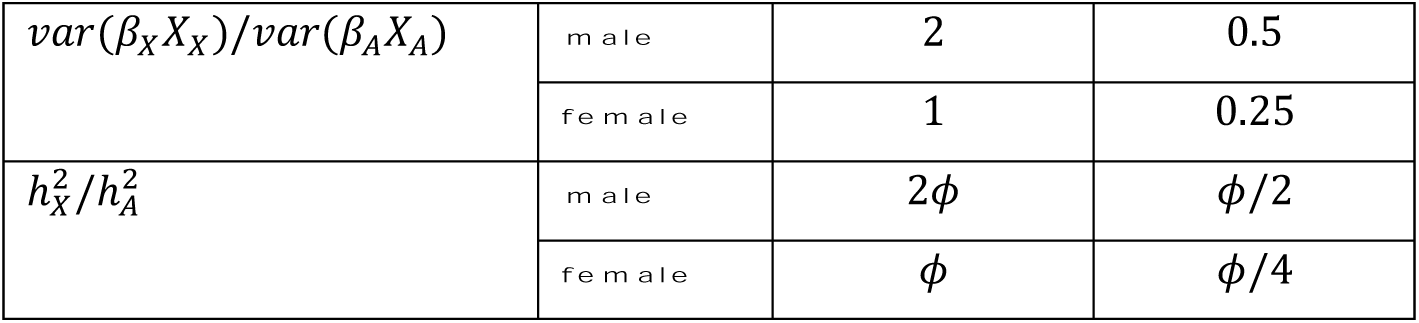
Expected relationship of quantities between sexes and between chrX and autosomes. *p*, the minor allele frequency, expected to be same for effect alleles in chrX and in autosomes. *var*_*m*_, *var*_*f*_, genotype variance of chrX in male and female analyses, respectively. *β*_*m*_, *β*_*f*_, male and female effects in chrX, respectively. *var*(*β*_*m*_*X*_*m*_), *var*(*β*_*f*_ *X*_*f*_), phenotypic variance explained by an X-linked variant with genotypes *X*_*m*_ and *X*_*f*_ in males and females, respectively. 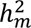, 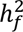, chrX heritabilities in females and males, respectively. *a*_*X*_, *a*_*A*_, active allele effects in chrX and autosomes, respectively. *β*_*X*_, *β*_*A*_, chrX and autosomal effects, respectively. *var*(*β*_*X*_ *X*_*X*_), *var*(*β*_*A*_*X*_*A*_), phenotypic variance explained by chrX and autosomal variants, respectively. 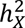, 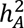, chrX and autosomal heritabilities, respectively. *ϕ*, the ratio of the number of variants contributing to heritability in chrX to that in autosomes. Here, *h*^2^ is defined as *var*(*βX*)/*var*(*Y*), where *var*(*Y*) is the total variance of the trait.

| Genotype variance of chrX variants ($var(X)$): | | |
| --- | --- | --- |
| $var_m = 4p(1 - p)$ | | |
| $var_f = 2p(1 - p)$ | | |
| Assuming full XCI and same active allele effect between sexes in the chrX ($a_m = a_f$): | | |
| $\beta_m, \beta_f$ – half of active allele effect size | | |
| $\beta_m = \beta_f = \frac{a_X}{2}$ | | |
| $var(\beta_m X_m)/var(\beta_f X_f) = 2$ | | |
| $h_m^2/h_f^2 = 2$ | | |
|  | X = AA | X = A |
| $a_X/a_A$ | 2 | 1 |
| $\beta_X/\beta_A$ | 1 | 0.5 |

|  |  |  |  |
| --- | --- | --- | --- |
| $var(\beta_X X_X)/var(\beta_A X_A)$ | male | 2 | 0.5 |
|  | female | 1 | 0.25 |
| $h_X^2/h_A^2$ | male | $2\phi$ | $\phi/2$ |
| | female | $\phi$ | $\phi/4$ |

The difference in ploidies between sexes is an inherent nature of chrX. Consequently, assuming full XCI and the same active allele effect in both sexes, the trait variance explained by a genetic locus (*β*^2^*var*(*X*)) is twice as large in males compared to females (Table 1). Thus, statistical power to detect a non-zero effect, which is an increasing function of the variance explained by the locus, is also larger in males than in females (assuming non-genetic variance of the trait is similar between the sexes). When performing a sex-specific analysis, the number of significant loci in male GWAS is expected to be larger than that in female GWAS in chrX given equal effect sizes, minor allele frequency (MAF) and sample sizes. For sex-combined GWAS, a common approach to carry out such analysis is a fixed-effect meta-analysis of sex-specific GWAS assuming *β*_*f*_ = *β*_*m*_ = *β*. For a chrX locus whose effect size differs between the sexes, this analysis has larger power to detect male-biased loci (|*β*_*m*_| > |*β*_*f*_ |) than female-biased loci (|*β*_*f*_| > |*β*_*m*_|).

To exemplify this power bias, we calculated the magnitude of the male or female effect size needed for 80% power to detect a SNP with MAF = 0.25 in chrX or in an autosome in a sex-combined GWAS with equal sample sizes of males and females (*n*_*m*_ = *n*_*f*_ = 180,000) at genome-wide significance level of 5 × 10^−8^. Under full XCI, where male-to-female effect size ratio is one, we were able to detect SNPs with effect size of ∼0.014 in chrX while a larger effect size of ∼0.017 was required in autosomes. If effect sizes differed between the sexes, we would be able to detect the sex-biased SNPs symmetrically in autosomes but with a bias favoring the male-biased SNPs in chrX in a sex-combined analysis (Figure S2). For instance, in a sex-combined chrX analysis, with a power of 80%, we can only detect a female-biased variants with twofold larger effects in females if the underlying female effect is as large as 0.021, while for a male-biased variants with twofold larger effects in males the underlying male effects need only be 0.017. A symmetric detection of sex-biased effect sizes in chrX under full XCI would require the sample size of males to be half of that of females to compensate for the doubled variance explained in males.

The escape from XCI is predicted to result in an increase in female effect compared to male effect at a locus level, and thus also result in an increased additive genetic variance in females^4^. Therefore, at a variant level, we can search for potential escape regions among those regions where effects are larger in females than in males. However, due to the power bias in the sex-combined GWAS, we have less power to detect loci with moderate degree of female bias than those with a similar degree of male bias. Thus, the estimated proportion of potential escape loci detected in a sex-combined analysis may be an underestimate because of the male-biased locus discovery in chrX. At a chromosome level, the escape from XCI may be small as it is reported to affect 15-30% genes^34^ and a previous study had found limited effect of escape when considering the contribution of all X-linked complex trait loci^4^.

### Genotype and phenotype data in UK Biobank

We used genotype datasets from UKB^27^ (release version 3 of imputed genotype data) for all analyses in the study. The details on genotyping, quality control, and imputation have been described previously^27^. Participants provided electronic signed consent at recruitment. Ethics approval for UKB was obtained from the North West Centre for Research Ethics Committee (11/NW/0382). All experiments were performed in accordance to relevant guidelines and regulations including the Declaration of Helsinki ethical principles for medical research. This study was run under UKB application number 22627.

The samples were included based on the following four criteria reported by the sample quality control file (“ukb_sqc_v2.txt.gz”) from UKB:

1. Not an outlier for heterozygosity and missing rates (the “het.missing.outliers” column);
2. Do not show putative sex chromosome aneuploidies (the “putative.sex.chromosome.aneuploidy” column);
3. Reported sex matches with inferred sex (the “Submitted.Gender” and “Inferred.Gender” column);
4. Were included in relatedness calculations (the “excluded.from.kinship.inference” column).

We restricted our analyses to unrelated White British, which were defined as those with a KING’s kinship coefficient lower than 0.04442. And we used the “in.white.British.ancestry.subset” column in the sample quality control file to define the White British. We removed individuals who had withdrawn from UKB by the time of this study.

All phenotypes were adjusted for males and females separately with appropriate covariates and inverse-normal transformed (Table S1), with values over six standard deviations from the mean removed as outliers prior to the normalization. For forced vital capacity, diastolic (DBP) and systolic blood pressures (SBP), the means were taken for individuals with repeated measures. For individuals on blood pressure medications at baseline measurement (UKB field ID: 6153 and 6177), 15 and 10 mmHg were added to their measured values of SBP and DBP, respectively. UKB includes measurements for 34 blood and urine biomarkers. In this study, rheumatoid factor, oestradiol, and microalbumin were excluded due to the large amount of missing data caused by the detection limits. For the remaining 31 biomarkers, we performed statin usage adjustment as described previously^35^. Briefly, we retrieved medication information from Treatment/medication (UKB field ID: 20003) and identified 1296 individuals who were not on statin during the initial visit (year 2006-2010) but were on statin during their first repeat visit (year 2012-13). A statin correction factor was calculated for each biomarker by taking the mean value of the ratio of on-statin measurement to pre-statin measurement. For 56,360 individuals who were taking statins upon enrollment, their biomarker measurements were divided by the statin correction factor to yield the adjusted values. The pre- and on-statin values were compared and only biomarkers showing a significant difference (*P*-value < 0.05, paired Wilcoxon rank-sum test) were adjusted with the statin correction factors (Table S1).

For non-biomarker quantitative traits, we included traits with estimated autosomal *h*^2^ ≥ 10% in sex-combined population in UKB SNP-Heritability Browser (see Web resources) that are available for both sexes. To avoid taking redundant traits (e.g. traits like impedance of left and right legs), we performed a hierarchical clustering and identified 15 clusters (Figure S3) based on the correlation of adjusted and normalized values between these traits in sex-combined population. Within each cluster, the trait with the highest *h*^2^ estimated in the UKB SNP-Heritability Browser was selected for GWAS. In addition, DBP and body mass index (BMI) were included for their medical relevance.

### Genome-wide association analyses in UKB

In all analyses, the non-PAR genotypes were coded as {0,2} in males and {0,1,2} in females. Both sex-specific and sex-combined GWAS were performed with BOLT-LMM v2.3.2^29^ for autosomes and chrX (both PAR and non-PAR). Directly genotyped variants (version 2) with MAF > 0.01 and missingness < 10% in autosomes and non-PAR chrX were used as the set of model SNPs in BOLT-LMM to estimate genetic relationship matrix and adjust the GWAS for confounding. GWAS statistics were calculated for imputed SNPs (version 3) with MAF > 0.001 and imputation quality > 0.7.

### Validation in FinnGen

The FinnGen data release 10 comprised of 430,897 genotyped Finnish individuals. The details on genotyping, quality control and imputation have been described previously^28^. Briefly, genotype imputation was performed with SISuV4.2 reference panel including 8556 high coverage whole genome sequenced Finnish individuals for autosomes and non-PAR of chrX. We performed sex-specific GWAS for height, BMI and weight with an average of 139,247 males and 154,408 females per trait using REGENIE v2.2.4 pipeline with similar covariates included as in the UKB GWAS (Table S1).

All participants in FinnGen provided informed consent for biobank research, based on the Finnish Biobank Act. Cohorts collected before the start of FinnGen (August 2017) were collected under study-specific consents and subsequently transferred to the Finnish biobanks following approval from Valvira, the National Supervisory Authority for Welfare and Health. Recruitment protocols followed the biobank protocols approved by Valvira. The Coordinating Ethics Committee of the Hospital District of Helsinki and Uusimaa approved the FinnGen study protocol Nr HUS/990/2017. The FinnGen study is approved by Finnish Institute for Health and Welfare (THL), approval number THL/2031/6.02.00/2017, amendments THL/1101/5.05.00/2017, THL/341/6.02.00/2018, THL/2222/6.02.00/2018, THL/283/6.02.00/2019, THL/1721/5.05.00/2019, Digital and population data service agency VRK43431/2017-3, VRK/6909/2018-3, VRK/4415/2019-3 the Social Insurance Institution (KELA) KELA 58/522/2017, KELA 131/522/2018, KELA 70/522/2019, KELA 98/522/2019, and Statistics Finland TK-53-1041-17. The Biobank Access Decisions for FinnGen samples and data utilized in FinnGen Data release 10 include: THL Biobank BB2017_55, BB2017_111, BB2018_19, BB_2018_34, BB_2018_67, BB2018_71, BB2019_7, BB2019_8, BB2019_26, BB2020_1, BB2021_65, Finnish Red Cross Blood Service Biobank 7.12.2017, Helsinki Biobank HUS/359/2017, HUS/248/2020, HUS/150/2022 §12, §13, §14, §15, §16, §17, §18, and §23, Auria Biobank AB17-5154 and amendment #1 (August 17 2020) and amendments BB_2021-0140, BB_2021-0156 (August 26 2021, Feb 2 2022), BB_2021-0169, BB_2021-0179, BB_2021-0161, AB20-5926 and amendment #1 (April 23 2020)and it’s modification (Sep 22 2021), Biobank Borealis of Northern Finland_2017_1013, 2021_5010, 2021_5018, 2021_5015, 2021_5023, 2021_5017, 2022_6001, Biobank of Eastern Finland 1186/2018 and amendment 22§/2020, 53§/2021, 13§/2022, 14§/2022, 15§/2022, Finnish Clinical Biobank Tampere MH0004 and amendments (21.02.2020 & 06.10.2020), §8/2021, §9/2022, §10/2022, §12/2022, §20/2022, §21/2022, §22/2022, §23/2022, Central Finland Biobank 1-2017, and Terveystalo Biobank STB 2018001 and amendment 25th Aug 2020, Finnish Hematological Registry and Clinical Biobank decision 18th June 2021, Arctic biobank P0844: ARC_2021_1001.

### Estimation of SNP-heritability and effect size distribution in chrX and autosomes

GENESIS^36^ is a likelihood-based approach for estimating effect size distribution and heritability and by default only analyzes autosomal summary statistics as only autosomal SNPs were included in the reference panel. To extend it to include the chrX, we extracted the tagging SNPs and calculated the corresponding LD-scores in chrX as described in ^36^. Briefly, we included HapMap3 SNPs with MAF ≥ 0.05 in the 1000 Genomes Project Phase 3 study^37^ of 489 individuals of European origin^38^ for PAR region and 256 females of European origin^38^ for non-PAR as our reference panel for chrX. Here, the tagging SNPs for a GWAS SNP were defined as those in the reference panel that were within 1 Mb distance and had an estimated LD coefficient (*r*^2^) with the GWAS marker above 0.1. We calculated the corresponding LD score for each GWAS marker by summing up the *r*^2^ for all tagging SNPs using 1000 Genomes Project data of both sexes for PAR and only females for non-PAR. We adjusted the LD score for bias as in ^36,39^. In total, 38,231 X-linked common variants (439 in PAR and 37,792 in non-PAR) were included in the reference panel for GENESIS analysis.

We analyzed the sex-specific summary statistics from GWAS for each trait with two-component model (referred as M2 in GENESIS) for the 38,231 common variants in chrX. We performed the analyses with ∼1.1 million common variants (MAF ≥ 0.05, excluding the Major Histocompatibility complex (MHC) region) in autosomes that was already included in GENESIS. GENESIS estimates the proportion of non-null effect variants (*π*_*c*_) and *h*^2^ explained per causal variant (*σ*^2^) that together define the effect size distribution. The SNP-heritability is defined as *h*^2^ = *M*_*c*_*σ*^2^ = *Mπ*_*c*_*σ*^2^, with *M*_*c*_ being the number of causal SNPs that is determined by *π*_*c*_ and *M*, the total number of HapMap 3 SNPs (38,231 in chrX and in 1,070,777 in autosomes).

For comparison, we estimated SNP-heritabilities of the 48 traits with LDSC^39^ using only autosomal sex-specific summary statistics. Precomputed LD scores of European individuals in 1000 Genomes Project were used as reference consisting of ∼1.2 million variants in autosomes^39^.

We tested whether 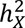 = 0 using the test statistic,

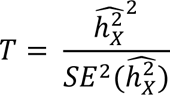

where 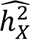 is the GENESIS estimates of SNP-heritabilities in chrX and *SE*(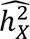) is the standard error (SE) of 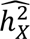. The test statistic follows a 𝜒^2^-distribution with one degree of freedom. The obtained *P*-values and false discovery rate (FDR) (using Benjamini-Hochberg procedure) are reported in Table S2.

### Estimation of X-chromosome influence (XI)

We estimated the contribution of chrX in complex trait genetics by defining the XI as

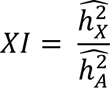

where 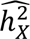 and 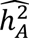 are the GENESIS estimates of SNP-heritabilities in chrX and autosomes respectively.

The SE of XI was estimated as

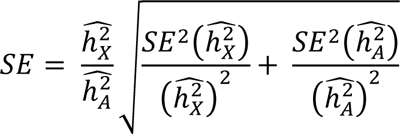

where *SE*(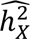) and *SE*(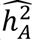) are the corresponding SEs of 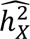 and 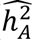 estimated with GENESIS.

We compared the observed XI to the XI predicted by the ratio (*ϕ*) of the number of variants in chrX to that in autosomes that contribute to heritability under two scenarios, X = AA and X = A (see Table 1). When the active allele effects of chrX are twofold compared to autosomes (X = AA), we expected the XI to be 2*ϕ* in males, *ϕ* in females, and 3*ϕ*/2 (between-sex mean) in the sex-combined population. When the active allele effects are equal between chrX and autosomes (X=A), we expected the XI to be *ϕ*/2 in males, *ϕ*/4 in females, and 3*ϕ*/8 in the sex-combined population.

We approximated *ϕ* ≈ 0.034 as the ratio of number of variants with MAF ≥ 0.01 in chrX to that in autosomes in the European population of the 1000 Genomes phase 3^37^. We also included the results with *ϕ* estimated based on the number of LD blocks present in chrX region and in autosomes for reference. The semi LD independent blocks were estimated using the LAVA partitioning algorithm^40^ using 263 female individuals of European ancestry of phase 3 of 1000 Genomes^37^, resulting in 71 LD blocks in non-PAR and 5 LD blocks in PAR (Table S7). We performed the partitioning with the same parameters as had earlier been used for partitioning the autosomes (excluding the MHC region) into 2479 LD blocks, that is, the default values of LAVA except that the minimum block size was set to 2500 as in^40^. Based on the number of LD blocks, *ϕ* ≈ 0.031.

### Identification of lead variants

Summary statistics of sex-specific GWAS were used to identify associated regions for each sex. For a SNP with a *P*-value (non-infinitesimal model) below 5 × 10^−8^, a region of 0.5 Mb around the SNP was defined as the association region. Overlapping regions were merged and considered as the same association signal. The variants with the smallest *P*-value within each region were considered as the lead variants. To compare effect sizes between autosomes and chrX, the effect size estimates and the corresponding standard errors of variants within non-PAR were multiplied by 2 in both male and female-specific analyses to estimate the per active allele effects. This was done since the functionally hemizygous variants in non-PAR were analyzed as diploid under the coding scheme used for chrX.

We performed conditional analysis on UKB sex-combined GWAS for each associated region with FINEMAP v1.4^41^. The analysis was performed with default settings but allowing for a maximum of 30 causal SNPs (--n-causal-snps 30) and the posterior probability of a causal configuration to be zero if the absolute correlation of two SNPs is above 0.9 (--corr-config 0.9). We used LD computed from UKB genotype data with LDstore v2.0 as recommended previously^42^.

### XCI scenarios analysis with sex-specific heritabilities

In theory, under full XCI (F-XCI), the 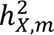 is expected to be twice that of the 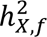 in chrX^4^ (Figure S1, Table 1). The absence of XCI (no XCI (N-XCI)) in females is expected to increase the female effect size twofold and hence result in male-to-female 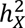 ratio of 0.5. Escape from XCI is expected to increase the 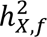, yet to a much smaller degree than N-XCI, as escape affects only a fraction of the chrX loci and typically in a manner where the expression from the inactive X remains partially suppressed. To derive a meaningful male-to-female 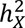 ratio for partial escape from XCI (E-XCI), we took the assumption of 25% of chrX loci undergoing escape. Further, following findings from gene expression studies^15^, where the expression from the inactive chrX is observed to be on average 33% of the expression from the active chrX, we assumed the effects from the inactive chrX to remain smaller than from the active chrX. To this end, we modeled the escape loci to follow approximately the relationship *β*_*f*_ = √2 · *β*_*m*_ (Figure S1), i.e., a ratio √2 ≈ 1.4 of female-to-male effects (here we analyzed effects under {0,1,2} and {0,2} coding scheme, but the same relationship can be assumed for per active allele effects, i.e. {0,0.5,1} and {0,1} coding scheme). Together these assumptions translate to a male-to-female 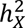 ratio for E-XCI at 1.75 (= 2*75% + 1*25%).

We applied the “linemodels” package^43^ to the sex-specific *h*^2^ estimates of 34 traits with nonzero 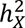 in both sexes. We clustered the traits into three groups that were represented by line models, whose slopes were set to 2 (F-XCI), 1.75 (E-XCI), 0.5 (N-XCI) for the chrX analysis and to 1(F-XCI), 0.875 (E-XCI), 0.25 (N-XCI) for the autosomal analysis assuming the same relationship between male and female genetic effects as in the chrX for the different XCI scenarios. For all models, the initial values for the scale parameters were set as the larger standard deviations of the *h*^2^ estimates across traits between male and female, the correlation parameters were fixed at 0.999, an equal prior probability across the models was assumed and the correlation of male and female *h*^2^ estimators was set to 0 because the samples were disjoint. The scale parameters were optimized in a two-step manner: first, we forced equal scales for all models by setting force.same.scales = TRUE in line.models.optimize() function; second, we used the optimized scale parameters and estimated proportions of models as initial values in line.models.optimize() function when we allowed different values for the scale parameters (force.same.scales = FALSE). Following the optimization of scale parameters for three models, we estimated the posterior probabilities in the three models separately for each trait with an equal prior probability assumed for each model. The analyses were performed separately for chrX and autosomal *h*^2^ estimates.

### Four-component sex bias mixture model of genome-wide variants

To detect moderate sex-biased effects of variants across the genome, we used a mixture model with four components: null effect (M0), female-biased effect (M1), equal effect (M2), and male-biased effect (M3). The mixture model was constructed and fit in STAN (version 2.21.0)^44^. The distribution of each component was formulated with sex-specific summary statistics with effect size 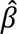 and its 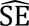 scaled multiplicatively by 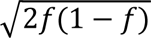, where *f* was MAF, so that, a priori, every variant explained similar phenotypic variance.

We denote by *β*_*m*_ and *β*_*f*_ the true male and female effects, respectively. Given the values of prior variance of the effect (*σ*^2^) and a parameter *α* > 1, the prior distributions of the components were

- M0: Null effect, *β*_*m*_ = *β*_*f*_ = 0;
- M1: Female-biased effect, *β*_*f*_ = *αβ*_*m*_, *β*_*m*_ ∼ *N*(0, *σ*^2^);
- M2: Equal effect, *β*_*m*_ = *β*_*f*_ ∼ *N*(0, *σ*^2^);
- M3: Male-biased effect, *β*_*m*_ = *αβ*_*f*_, *β*_*f*_ ∼ *N*(0, *σ*^2^).

Thus, *σ*^2^ was assumed the same for each non-null component and its prior distribution was Uniform(0,1) (see Supplemental Note for the choice of this prior). Additionally, the model includes parameter vector *π* = (*π*_0_, *π*_1_, *π*_2_, *π*_3_), where *π*_*k*_ describes the proportion of variants belonging to component *k*. For *π*, we used Dirichlet(1/4, 1/4, 1/4, 1/4) distribution as the prior to not favor any component a priori.

This model together with Normally distributed effect size estimates with known standard errors leads to the following marginal distribution for the observed data (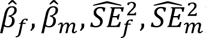):

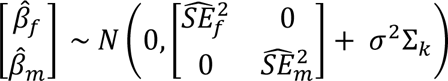

where for each non-null model the correlation between 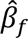 and 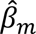 was assumed 1:

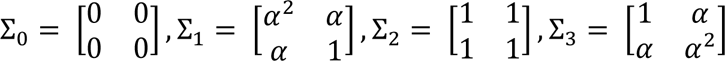

In our analyses, we set *α* = √2 because, in the female-biased component, as we expected the effect of escape from XCI would result only in a moderate, clearly less than twofold increase in female effect compared to male effect.

Using this model, we estimated *σ*^2^, the variance of the non-null effects, and *π*_*k*_′*s*, the proportions of variants belonging to each component.

To identify variants with sex-biased effects, the estimated parameters were used to calculate a probability for variants to be assigned to a given component. For variant *i*, the probability *p*_*i*,*k*_ for it to be in component *k* is ^45^:

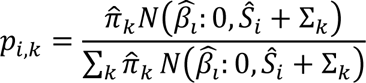

Where 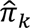 is the posterior mean of proportion estimated by the model and 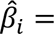 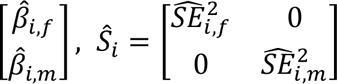. Variants were assigned to *k* component if the posterior probability *p*_*i*,*k*_ > 0.8; otherwise, they were left “uncategorized”.

We focused on variants outside of the MHC region and excluded variants with missingness > 1% and MAF < 0.01 in both sexes in both autosomes and chrX, and Hardy-Weinberg disequilibrium test *P*-value < 10^−7^ followed by LD pruning with PLINK 1.9^32^ using “--indep-pairwise 50 5 0.1” with genotype data from both sexes for autosomes and PAR and genotype data from females for non-PAR. This resulted in 4380 variants in chrX and 152,091 variants in autosomes. For chrX, the analyses were performed using all 4380 variants in chrX. For autosomes, six variants were sampled per LD group (total 1693 LD groups) estimated in Europeans^46^ resulting in 10,158 variants to allow efficient estimation of the parameters. All models were run with 4 chains, using 2000 warm-up and 4000 total iterations. Convergence was assessed using Rhat, which measures consistency of chains, and traits with parameter fits with Rhat greater than 1.01 were excluded.

## Results

### Contribution of chrX to complex traits

We first performed sex-specific GWAS, using both autosomal and chrX, with BOLT-LMM^29^ in UKB for 48 quantitative traits (see details of traits selection in Material and methods and trait information in Table S1). For validation purposes, we conducted similar association analyses for three of the UKB traits (height, BMI, and weight) in FinnGen^28^ (release 10).

We asked how much the additive genetic effects in chrX contribute to quantitative trait variation in the overall population. To this end, we estimated the sex-combined 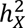 as the average of the male and female-specific estimates from GENESIS^36^, which we extended to allow the inclusion of chrX (see Material and methods). Sex specific data was used to avoid the power biases impacting the analyses of chrX variation in sex-combined data (see Material and methods). For comparison, the same method was applied to autosomal data. We observed a clear role of chrX variation in most complex traits as 45 out of the 48 analyzed UKB traits showed 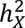 estimates significantly different from zero (FDR < 0.05), with height displaying the highest 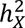 (UKB: 2.89% (SE = 0.30%); FinnGen: 3.04% (SE = 0.31%)) (Table S2).

We next compared the estimated 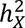 to the corresponding 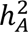 to understand the relative importance of chrX variation in complex traits. Across all the analyzed traits, 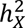 tracked with 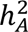 (Pearson *r* = 0.88; Figure 1A) indicating the contribution from chrX to a complex trait is typically proportional to the autosomal contribution. To further quantify the role of chrX in quantitative trait variation, we defined the chrX influence (XI) as the ratio of 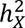 to 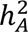 and calculated this quantity separately for each trait. We observed a median XI of 0.03 (Figure 1A) suggesting that, in the overall population, chrX contributes to complex trait heritability an additional 3% of the contribution of autosomes.

**Figure 1.**
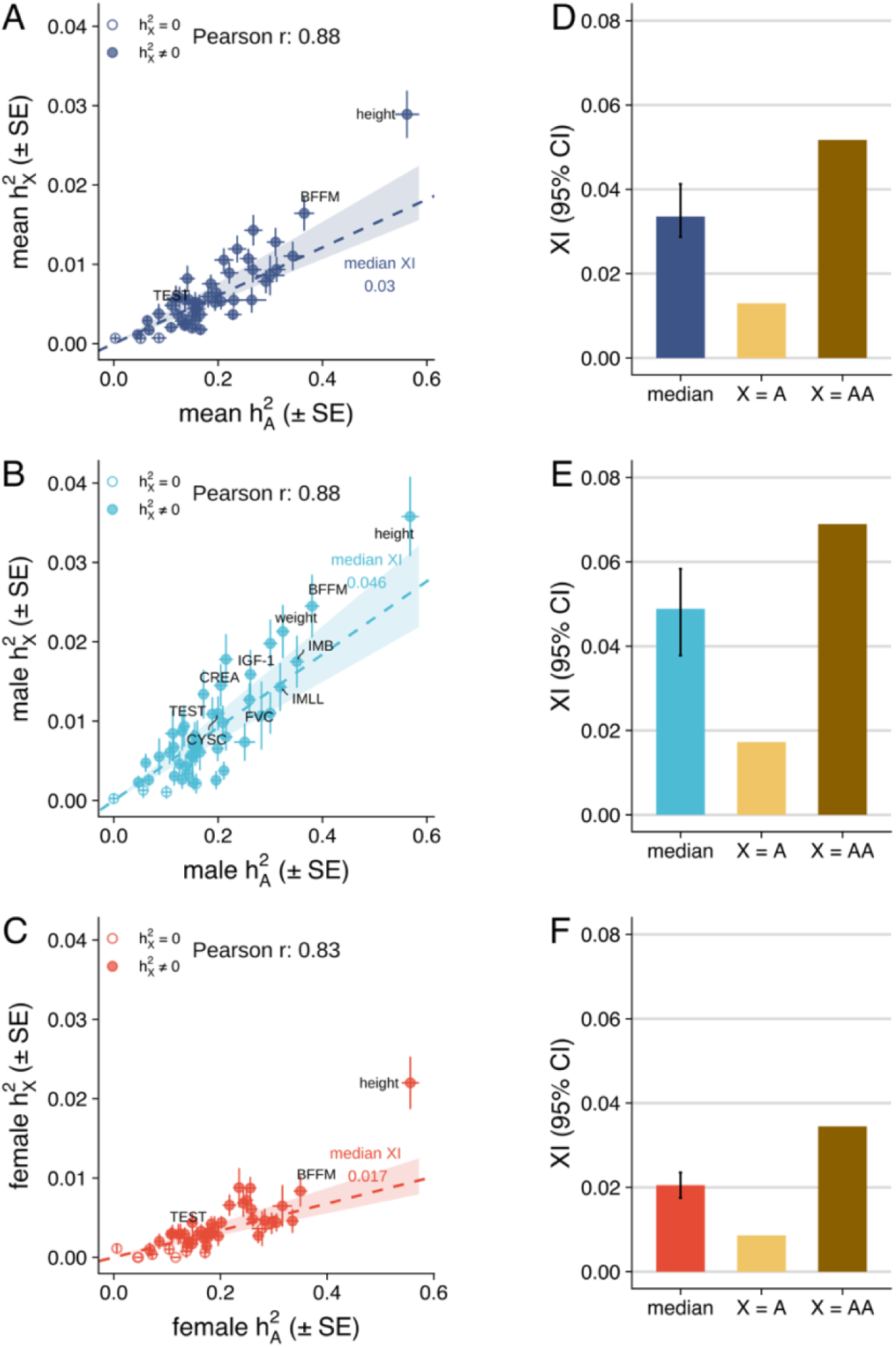
The contribution of chrX to complex trait genetics. 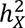 versus 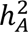(A) in the overall population estimated as the average of sexes, (B) in males, and (C) in females. Dashed lines indicate the median XI, i.e. 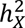 /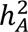, with shaded area indicating bootstrap 95% CI of the median. For 35 traits with nonzero 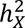 in both sexes, the median XI (bootstrap 95% CI) is compared to the expected XI derived based on the proportion of common variants in chrX and assuming the genetic effect of one active X is equal to one copy of an autosome (X = A) or to a pair of an autosome (X = AA) (D) in the overall population, (E) in males, and (F) in females. Numeric results are reported in Table S2. Abbreviations: whole body fat-free mass (BFFM), impedance of body (IMB), insulin-like growth factor 1 (IGF-1), impedance of leg, left (IMLL), forced vital capacity (FVC), testosterone (TEST), cystatin C (CYSC).

Given the unique sex-dependent biology of chrX, we assessed how chrX contributes to complex trait variation differently between sexes by comparing the sex-specific *h*^2^ estimates. As expected, given the chrX dosage difference between the sexes and XCI in females (see Material and methods), we observed, in general, higher 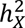 in males compared to females (mean 0.88% (SD = 0.69%) versus 0.37% (SD = 0.69%); *P*-value = 3.54 × 10^−1^^1^, paired Wilcoxon rank-sum test), consistent with earlier reports^4,5^. This higher 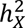 in males is reflected also in the 2.3-fold greater number of X-linked genome-wide significant loci in males compared to females (Figure S4 and Supplemental Note). In contrast, we observed no systematic sex difference in 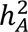 (*P*-value = 0.89, paired Wilcoxon rank-sum test), although a few traits (5/48), namely testosterone, DBP, urate, insulin growth factor-1 and waist-to-hip ratio (WHR), showed a significant sex difference in 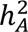 (FDR < 0.05) as reported previously^45,47–49^ (Table S2).

As expected, given the above results, a clear sex difference was also observed in XI, with a consistent pattern of greater XI in males compared to females (Figures 1B and 1C; median 0.046 versus 0.017; *P*-value = 3.91 × 10^−1^^3^, paired Wilcoxon rank-sum test). Interestingly, DBP and SBP were exceptions to this pattern, with greater XI in females than in males (DBP: 0.018 (SE = 0.0036) versus 0.013 (SE = 0.0078); SBP: 0.029 (SE = 0.0066) versus 0.018 (SE = 0.0053)). Together, however, these observations exemplify the greater relative importance of the chrX variation in males compared to females arising from the impact of the ploidy difference and chromosome-wide inactivation in females.

### Interpreting XI through the lens of dosage compensation

To provide insights into the potential dosage compensation between chrX and autosomes, we compared the above XI results to theoretical expectations of chrX-to-autosomes relationship under two scenarios. In the first scenario (X = AA), one active chrX is equivalent to a pair of autosomes, with twofold active allele effects in chrX compared to autosomes. In the second scenario (X = A), one active chrX is equivalent to a single autosome, with equal active allele effects between chrX and autosomes (see Table 1 and Material and methods). For these theoretical expectations, we assumed that the allelic effects are small and uniformly distributed along the genome, and the expected values were computed based on the proportion of common variants (chrX contains ∼3.4% of the number of common variants in autosomes (see Material and methods)).

Focusing on the 35 traits with nonzero 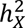 in both sexes, we observed that the medians of XI differed significantly (*P*-value < 0.05 based on bootstrap 95% CI) from the expected values under both scenarios (Figures 1D-F). In the overall population, the observed median XI (0.034, bootstrap 95% CI 0.029 – 0.041) was 1.5-fold lower than expected under X = AA (0.052) and 2.6-fold greater than expected under X = A (0.013) (Figure 1D). A similar degree of difference was observed in the sex-specific data (Figures 1E and 1F). Similar patterns were also observed with expected values derived based on the number of LD blocks of which chrX contains 3.1% of those in the autosomes (Figure S5).

Dissection of *h*^2^ into the proportion of causal variants (*π*_*c*_, estimates polygenicity) and per-SNP-*h*^2^ (*σ*^2^, estimates the magnitude of nonzero effects) suggested the mismatch between the observed XI and assumptions under the dosage compensation models arises from the effect sizes rather than from systematic differences in the polygenicity between chrX and the autosomes. While the estimated polygenicity varied greatly between chrX and autosomes (ratio of *π*_*c*_ from chrX and autosomes ranges between 0.3-4 for the 35 traits with nonzero 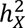 in both sexes) (Figure S6), the observed medians of 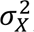/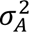 were again significantly lower than expected under X = AA (1.5 and 2.4-fold smaller than the expected in males and females, respectively) and higher than expected under X = A (2.5 and 1.7-fold greater than the expected in males and females, respectively) (Figure S7), closely mirroring the results from the XI comparisons. As heritability is a function of the squared effect size, the observed enriched per-SNP-*h*^2^ in chrX under X = A translate to a median of 1.6 and 1.3-fold larger per active allele effects in chrX than in autosomes in males and females, respectively.

These observations therefore suggest that, although there is only a single active copy of chrX in both sexes owing to the hemizygosity of men and XCI in females, the single active copy of chrX contributes to complex trait heritability more than a single autosomal copy with similar amount of genetic variation but less than two such autosomal copies.

### Comparison of active allelic effects between chrX and autosomes

To formally test for the differences in effect sizes, as suggested by the above results, we assessed the chrX-to-autosome differences in allelic effect estimates across the studied traits. To this end, we compared the effect sizes per active allele (see Material and methods) of lead variants in chrX (*a*_*X*_) to those of autosomes (*a*_*A*_) identified from the male and female GWAS, where the males were down-sampled by half in chrX association analyses to achieve similar power as in the autosomes (see Material and methods). We observed that the medians of *a*_*X*_ were 1.9 and 1.8-fold higher in chrX than that of *a*_*A*_ in males and females, respectively (0.054 versus 0.028 in males, 0.048 versus 0.026 in females; *P*-value = 1.43 × 10^−2^^8^ and 2.13 × 10^−5^ for males and females, respectively, Wilcoxon rank-sum test) (Figure 2A), a pattern not influenced by pleiotropic loci (Figure S8A). To eliminate the effect of the Winner’s curse, we further compared the effects estimated using FinnGen data for the height, BMI and weight associated variants identified from the UKB data. Although limited by the numbers of variants in chrX, the difference between the chrX and autosomes remained, with the medians of *a*_*X*_ being 1.6 and 1.7-fold higher than the medians of *a*_*A*_, in males and females, respectively (Figure 2B).

**Figure 2.**
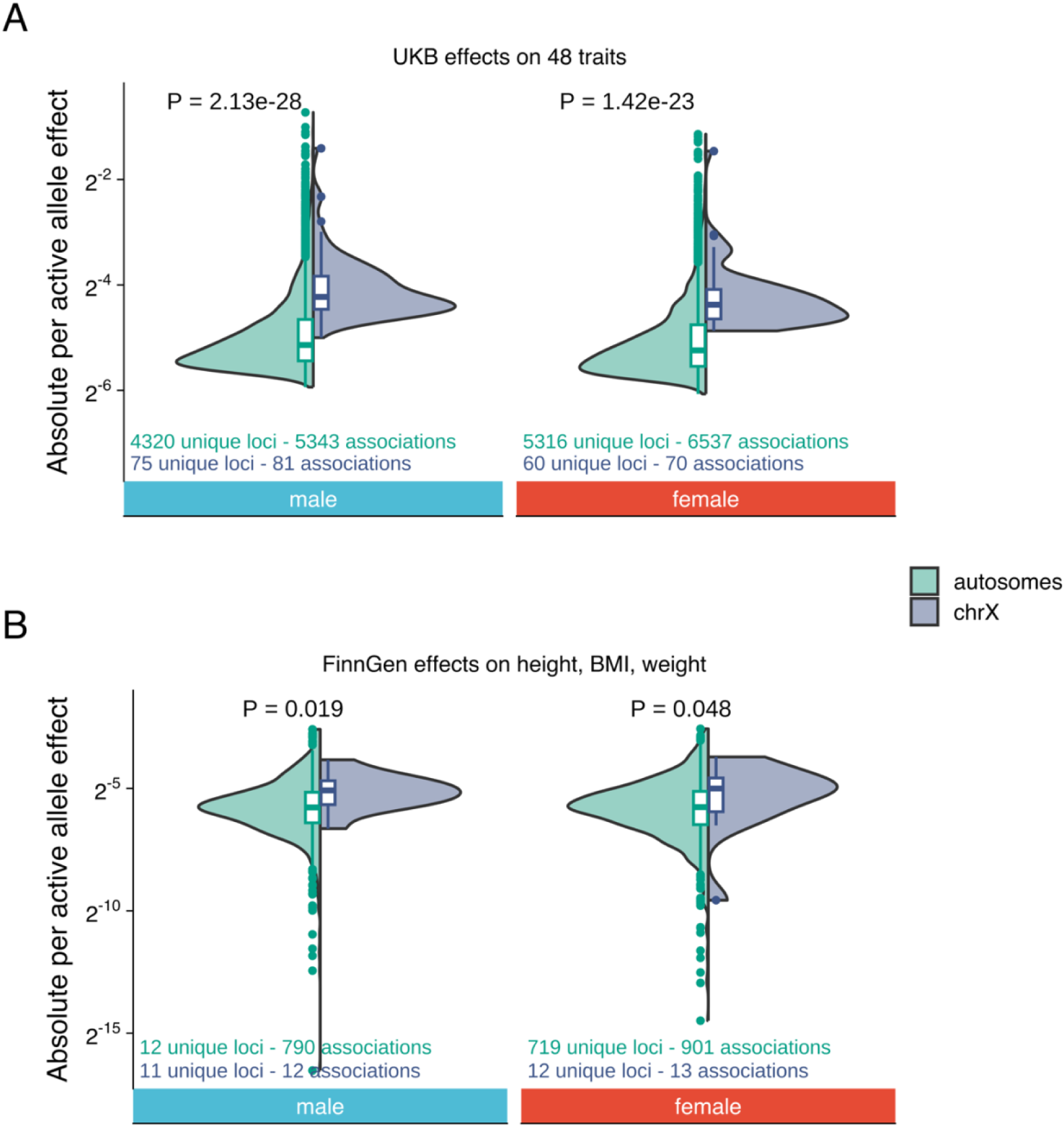
Comparison of per active allele effects between chrX and autosomes. Comparison of per active allele effect size (*a*) between autosomes and chrX for sex-specific trait-associated variants identified in UKB (A) with *a* estimated in UKB for all 48 traits and (B) with *a* estimated in FinnGen for height, BMI and weight. The male GWAS in non-PAR has been down-sampled by half to have similar statistical power as in autosomal GWAS. *P*-values for the difference between autosomes and chrX (Wilcox rank-sum test) are indicated on the top. Numerical values are reported in Table S6.

We assessed if these observations were explained by differences in MAF, functional consequences, or pathogenicity of variants between chrX and autosomes. We observed chrX overall has slightly higher MAF, less regulatory and coding regions and less pathogenic variants relative to autosomes. However, we observed systematically larger *a*_*X*_ than *a*_*A*_ independent of variant frequency or consequence. An exception was the most constrained regions, in which variants are rare in chrX and an upper bound may be imposed on the active allele effects by negative selection (Figures S9—11, Supplemental Note).

Taken together, these observations suggest that, overall, common variants in chrX have larger active allele effects compared to the autosomes, which likely explains the higher XI compared to the expected under X = A that only one functional copy of chrX is present in each cell.

### Insight into XCI escape through sex-specific heritability comparison

Our earlier results of 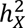 indicated a clear sex difference consistent with the presence of XCI (Figure 1B-C). To further understand the completeness of XCI through complex trait genetics, we compared how different XCI scenarios explain the observed relationship of male and female 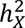. To this end, we applied a Bayesian approach^43^ to cluster the traits to the three XCI models: full XCI (F-XCI), no XCI (N-XCI) and escape from XCI (E-XCI) that accounts for a scenario where 25% of the chrX loci partially escape from XCI (see Material and methods).

Using a posterior probability threshold of 0.80, for most of the traits, we were unable to distinguish between the F-XCI model (male-to-female 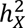 ratio = 2) and the E-XCI (male-to-female 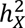 ratio = 1.75) model (Figure 3A). For instance, forced vital capacity had F-XCI posterior probability of 0.41 and E-XCI posterior probability of 0.59. However, DBP and SBP clustered to the N-XCI model with posterior probability of 0.98 (Figure 3A). For these blood pressure traits, we also observed larger 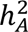 in females than in males (Figure 3B), as reported previously^50^, suggesting the sex differences in 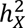 are unlikely to be explained by the lack of or escape from XCI alone but may be attributable to other factors such as hormonal influences^51^. The only trait that was best explained by the partial escape from XCI model was height (E-XCI posterior probability = 0.99) (Figure 3A), a trait that we found highly heritable in chrX (Figure 1A-C). Similar sex difference in 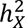 for height was also detected in FinnGen (E-XCI posterior probability = 1.00) (Figure 3A) but no sex difference was seen in autosomal data for height in either of the data sets (Figure 3B). The finding of female-enriched 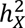 points to the potential role of chrX loci escaping from XCI in human height.

**Figure 3.**
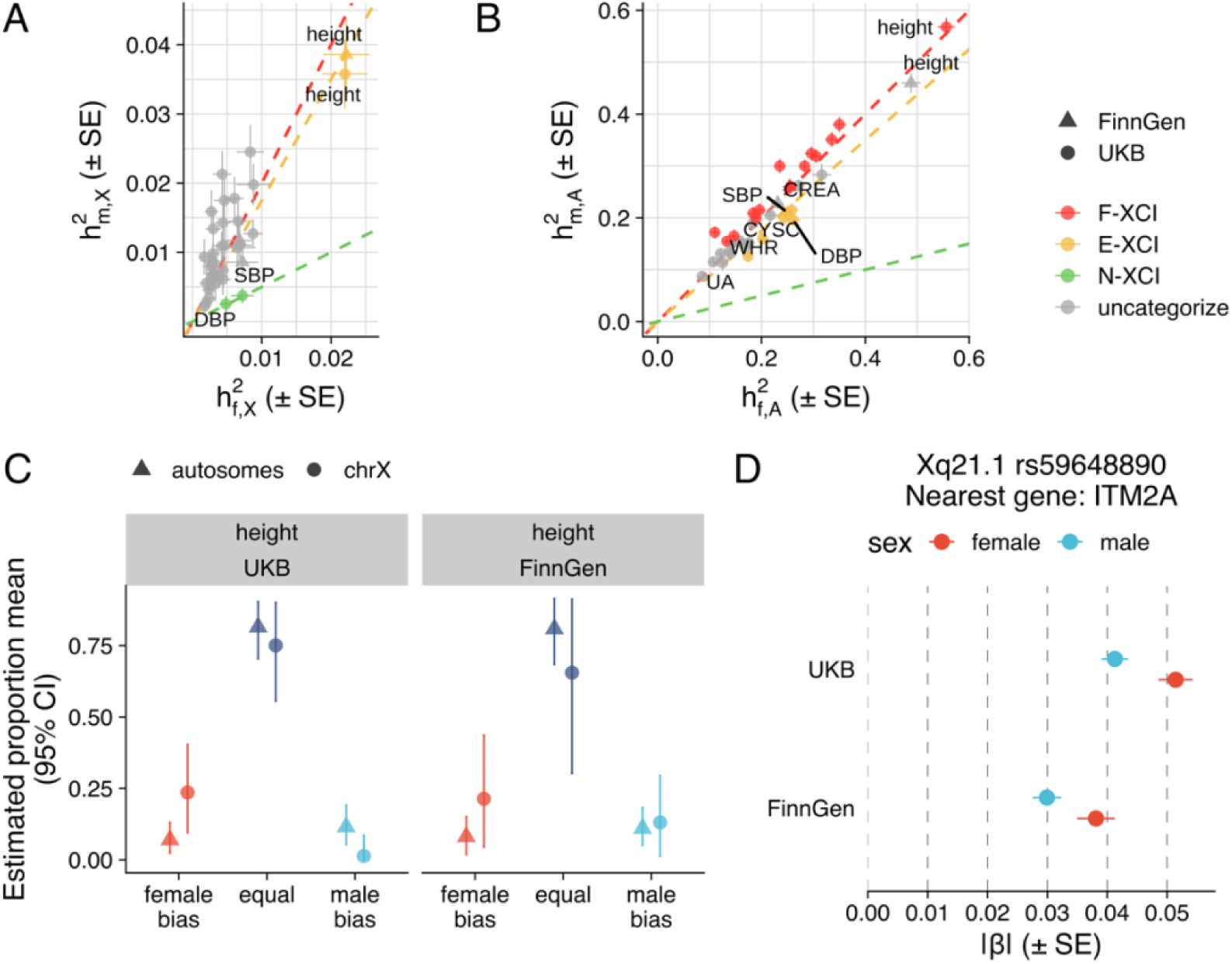
Comparison of male and female *h*^2^ and estimated SNP effects. Comparison of male and female *h*^2^ in (A) chrX and (B) autosomes for each trait and clustering based on theoretical XCI scenarios. The red, yellow and green dashed lines indicate expectation under full XCI (F-XCI), 25% escape from XCI (E-XCI), and no XCI (N-XCI), respectively. Colored points belong to the cluster with probability > 0.80. Numeric values are reported in Table S8. (C) Estimated proportion of variants with female-biased, equal and male-biased effect on height in UKB and FinnGen. Numeric values are reported in Tables S9. (D) Sex-specific effects of rs59648890 variant near *ITM2A* (integral membrane protein 2A) on height in UKB and FinnGen. Abbreviations: diastolic blood pressure (DBP), systolic blood pressure (SBP), creatinine (CREA), urate (UA), waist-to-hip ratio (WHR), cystatin C (CYSC).

### Comparison of sex-biased effects between chrX and autosomes

To further elucidate the potential sex-dependent genetic architecture in chrX, we asked if the proportions of variants with either female-biased or male-biased effects differ between chrX and autosomes. To this end, we applied a mixture model to sex-specific summary statistics to estimate the proportion of genetic effects with female or male bias (see Material and methods). The mixture model contains four components to capture the following types of variants: no effect on the trait in either sex, equal non-null effects in males and females, female-biased effects (|*β*_*f*_ | > |*β*_*m*_|), and male-biased (|*β*_*m*_| > |*β*_*f*_ |) effects.

Focusing on the proportions of variants with non-null effects, we observed that, compared to autosomes, there are proportionally fewer associations with equal genetic effects between sexes in chrX (*P*-value = 9.97 × 10^−8^, paired Wilcoxon rank-sum test of point estimates; Figure S12), pointing to unique sex-biased characteristics of chrX. This result was driven by a greater fraction in male-biased effects (*P*-value = 4.46 × 10^−6^, paired Wilcoxon rank-sum test of point estimates) rather than female-biased effects (*P*-value = 0.07, paired Wilcoxon rank-sum test of point estimates) in chrX. The enrichment of male-biased effects may be largely attributable to the pleotropic male-specific effects of regions in chrX associated with testosterone, a trait known for enriched male-specific effects in chrX^45,52^ (see Supplemental Note). Interestingly, for WHR, a trait known for largely female-biased genetic effects in the autosomes^53^, the majority of non-null variants were expectedly estimated to be female-biased in autosomes (78.8% (95% CI: 51.1% – 98.4%)), however, in chrX, a considerably smaller fraction, 16.8% (95% CI: 0 – 67.0%) of non-null variants, were estimated be in the female-biased component (see Supplemental Note).

Echoing the results of XCI analysis on male and female *h*^2^ comparison on height, we observed a greater proportion of non-null variants with female-biased effects in chrX compared to autosomes for height (23.6% (95% CI: 9.57 – 40.4%) versus 6.99% (95% CI: 2.35 – 13.1%)) (Figure 3D). To ensure this was not due to sex differential participation bias in the volunteer-based UKB^54^, we performed the same analysis in FinnGen, where the participation is more passive. We observed the same pattern of an enrichment of female-biased effects among the non-null variants in chrX (21.4% (95% CI: 4.47 – 43.7%) versus 8.09% (95% CI: 1.86 – 15.2%)) (Figure 3D).

To pinpoint individual loci driving the observed female bias in chrX for height, we computed the posterior probability of each component for the genome-wide significant height-associated variants identified in sex-combined conditional analysis (n = 73). We identified eight lead variants as female-biased (posterior probability > 0.80), however, only one variant was replicated in FinnGen as a female-biased variant (Figure S13A). This highlights the poor consistency of sex differences across biobanks despite the highly reproducible genetic effects (Figures S13 and S14, Supplemental Note). The replicated variant, rs59648890, locates 33 kb upstream of *ITM2A* (integral membrane protein 2A, a gene involving in cartilage development) (Figure 3E), confirming our earlier findings in a smaller Finnish sample^7^. Supporting the SNP level finding, the male-to-female ratio of local *h*^2^ at the LD block containing the *ITM2A* region (X: 77844781-80093260) was also smaller than expected under F-XCI at 2 (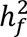= 0.41% and 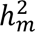 = 0.53% in UKB; 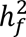 = 0.55% and 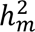 = 0.98% in FinnGen). The moderate female-biased effect (1.2 times greater in females than in males in UKB and 1.3 times in FinnGen) at the locus is aligned with partial escape from XCI.

## Discussion

ChrX has remained understudied in GWAS largely owing to the distinct challenges it poses for the analysis and interpretation of genetic associations. We set out to provide a thorough understanding on how the chromosome and its unique biology contribute to complex traits. Through analyzing large-scale biobank data across a broad panel of quantitative traits, we demonstrated that chrX hosts complex trait heritability and loci proportional to the contribution of autosomes. Our findings further support the presence of near-full XCI^4,5^, the dosage compensation between XY males and XX females, and highlight the relevance of this process for the sex-specific contributions of the chromosome for complex traits. Our results also mirror the dosage compensation mechanism between chrX and autosomes through X-upregulation proposed by Ohno^11^, whereby the contributions between chrX and autosomes are balanced through systematically larger chrX effect sizes per a single active copy of the chromosme.

Across the 48 complex traits investigated, we found that, in the overall population, the chrX *h*^2^ equaled to approximately an 3% of the autosomal *h*^2^. As such, the contribution of chrX to complex trait variation is typically less than that of the chromosome size, but is in line with the proportion of genetic variants in the chromosome, which is smaller than that of a similar sized autosome.

The chrX-to-autosome *h*^2^ ratio can also be interpreted in the light of dosage compensation where, due to XCI, only a single copy of chrX is active in each sex. The observed 3% contribution of chrX rather suggests a greater role than expected under X = A, but smaller than expected under X = AA. We attribute this finding to the systematically larger, but less than twofold, active allele effects in chrX compared to autosomes. Although previous studies have similarly implicated differences in effect sizes between chrX and autosomes, the differences in allele coding practices have resulted in mixed interpretations^4,26^. As we observed the effect size differences in both sexes, the result is not attributable to the hemizygosity of males or escape from XCI in females, neither is it due to the differences in allele frequencies nor variant constraint between common variants in chrX and autosomes. Rather, the finding appears to parallel Ohno’s dosage compensation hypothesis on a global transcriptional X-upregulation to account for the dosage difference between a single active chrX and pairs of autosomes. While less studied than XCI, X-upregulation has been shown to be present but partial through gene expression analyses^18–23^, proposed to occur through more frequent transcriptional bursting from chrX^22^. Interestingly, our estimate of the active allele effect size difference in complex trait genetic associations (∼1.6-fold) is close to the estimated degree of transcriptional X-upregulation (∼1.4-fold)^22^. However, whether these two phenomena share a mechanisitic basis warrants further investigations.

In the sex-specific analyses, we found that the XI has a clear sex difference, with on average doubled contribution to phenotypic variation in males, consistent with the joint effects of the male hemizygosity and XCI. These findings echo observations from other complex traits^4,5^ and align with the established male preponderance for X-linked disorders where random XCI in females confers protection. Although we find chrX has a role in many complex traits, the sex difference in chrX heritability due to XCI is unlikely to be reflected significantly in the genome-wide trait heritability estimates in the current sample sizes owing to chrX typically contributing only a few percentages of the overall heritability. Further, it follows, given the greater role of chrX variation in males, that additive effects in chrX are unlikely to explain female biases in complex phenotypes.

We additionally examine the role of escape from XCI in complex traits, a region-specific phenomenon in the chromosome, shown to result in sex-biased gene expression^15–17,55,56^. Altough genetic data from chrX could theoritically be used to identify escape regions and traits being impacted, the anticipated subtle changes on genetic effects due to escape render such assessments highly challenging even in biobank-scale datasets. Building on prior gene expression studies on partial escape^15–17^, we expected ∼25% of chrX loci may escape with female-to-male effect ratio at ∼1.4. Similar degree of female bias (female-to-male effect ratio at 1.5) was recently found for a locus in *PRKX*, a known escapee, in lymphocyte count association^25^. Following the typically modest *h*^2^ and generally small genetic effects, we find plausible evidence consistent with partical escape only in height, a trait where we had the largest power for the assessment. After replication, we could pinpoint plausibly the *ITM2A* locus not subject to full XCI. The *ITM2A* locus has been reported previously associated with height but evidence for escape interpreted differently^4,7^. Further validation and mechanistic dissection of this locus is warranted. While it is possible that even larger sample sizes will allow for more detailed quantification of escape in complex traits, the clear gene-specific differences related to escape seen at the transcriptome level can also be buffered through other processes before reaching more distal phenotypes. Overall, it is possible the proposed contribution of escape from XCI to phenotypic sex differences acts via other mechanisms than through direct locus-specific effects on phenotypes.

In the light of these findings, we propose that the two dosage compensation mechanisms possibly act in concert to optimally balance the role of chrX in the population. Owing to nearly full XCI, the per-SNP-*h*^2^ is about twice as high in males as in females, and resides in between in the overall population. Given this, under equal active allele effects between chrX and autosomes, the per-SNP-*h*^2^ in chrX would remain at relatively low level compared to autosomes in both sexes and so the contribution of chrX to complex traits would be much lower than expected by the amount of genetic variation in chrX (Figure 4). Twofold active allele effects in chrX would, on the other hand, increase the per-SNP-*h*^2^ to a high level in males, where a possible upper bound may be imposed by negative slection on the hemizygous chrX in males. A “partial upregulation” of chrX, with 1.6-fold larger active allele effects in chrX compared to autosomes, a scenario closely matching our observations, would balance out the sex difference in per-SNP-*h*^2^ in the overall population, resulting in comparable per-SNP-*h*^2^ between chrX and autosomes and the contribution of chrX to complex trait genetics in par with to that of the autosomes (Figure 4).

**Figure 4.**
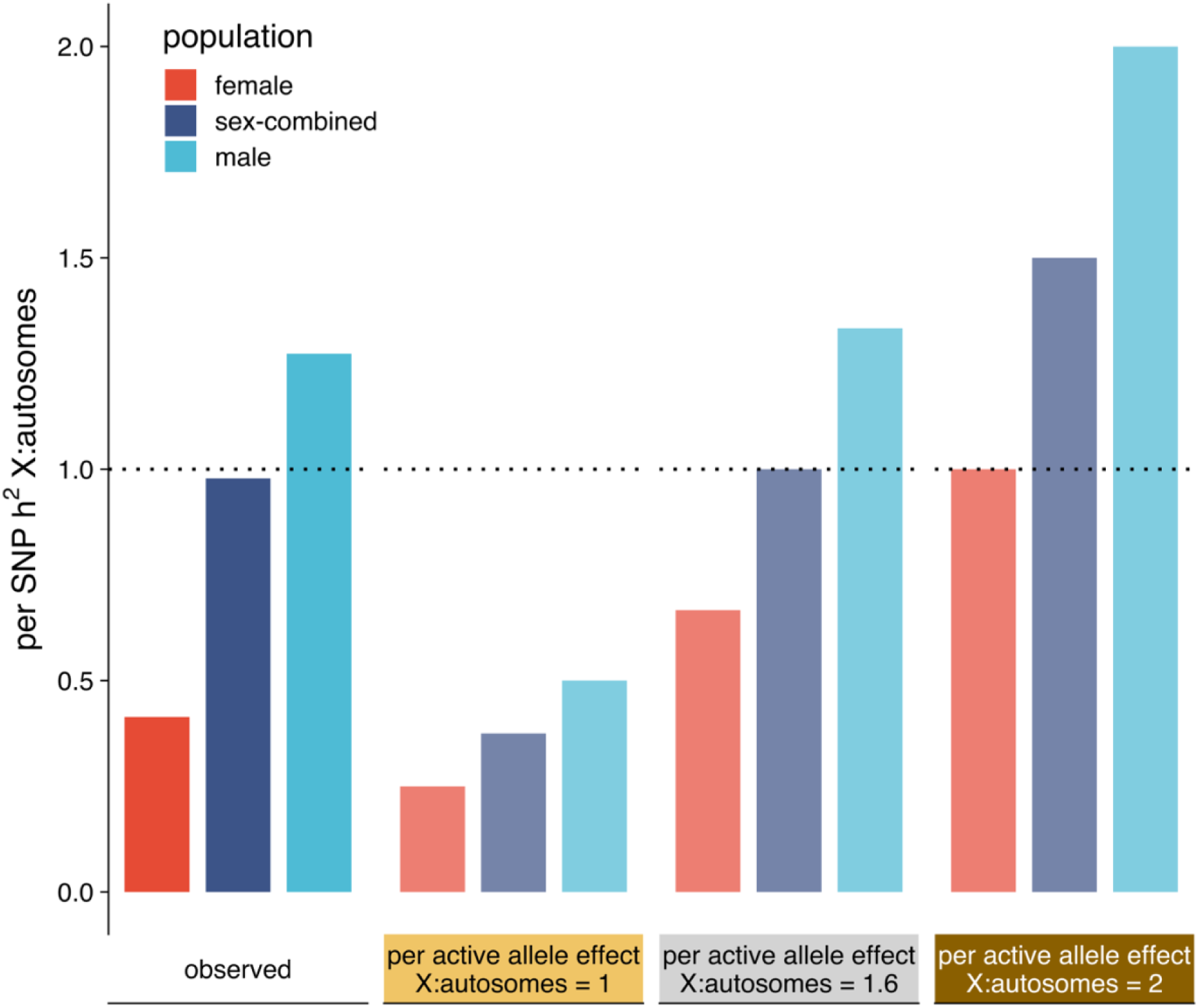
Model of how XI is optimally balanced in the population. Illustration of chrX-to-autosomes per-SNP-*h*^2^ ratio in male, female, and overall populations assuming full XCI for observed data (medians) and for three theoretical chrX-to-autosomes per active allele effect ratios. The observed per-SNP-*h*^2^ in the sex-combined population is estimated as the mean across males and females. The dotted line indicates when the per-SNP-*h*^2^ is equal between chrX and autosomes in the sex-combined population.

Our assessments in this study focused on the role of additive genetic variation in chrX. While additive effects are the primary mode of heritability in autosomes^57^, the unique characteristics of chrX can make other types of effects more relevant. For instance, X-linked deleterious alleles that are known to play a role in rare diseases often affect females in a recessive manner. Further, skewed XCI, that changes the heterozygous dosage in females to homozygous, may be particularly relevant for higher impact variation, and it has been observed at higher frequencies among individuals with autoimmune diseases^58,59^. Also, our assessments included only quantitative traits, and different dynamics may be expected when extending chrX analyses to complex diseases.

Our primary studies were performed in UKB, which is a volunteer-based study with evidence for sex-differential participantion bias in autosomes^54^. Thus, where possible, we set to validate our findings in FinnGen, a data set with more passive participation design. While we observed same female-biased pattern on the chromosome level for height, the sex-biased effects for individual loci were poorly replicated across biobanks, highlighting the broader challenges associated with the detection of gene-by-environment interactions^60,61^.

As most GWAS tools nowadays support the analysis of chrX, the inclusion of chrX, as shown in our study, offers a possibility to uncover new biology and trait *h*^2^. Although the underlying assumptions regarding dosage compensation in chrX analyses may not be highly relevant for locus discovery, these matter for the intepretation of the relationship of male to female and chrX to autosomes effects (see Material and methods). Further, while sex differences in complex trait genetic architecture are typically modest in chrX, one should nevertheless be aware of the power difference between the sexes that bias the detection towards male-biased effects when the variant selection is based on a significance threshold. The potential effects of escape, although shown here to be limited at the currently available sample sizes, may become evident as sample sizes in GWAS continue to grow.

Taken together, our work shows that in addition to providing new complex trait associations, GWAS data on chrX provides possibilities to delve in the unique biology of this chromosome.

## Acknowledgements

The research has been conducted using the UK Biobank Resource under application number 22627. The FinnGen project is funded by two grants from Business Finland (HUS 4685/31/2016 and UH 4386/31/2016) and the following industry partners: AbbVie Inc., AstraZeneca UK Ltd, Biogen MA Inc., Bristol Myers Squibb (and Celgene Corporation & Celgene International II Sàrl), Genentech Inc., Merck Sharp & Dohme LCC, Pfizer Inc., GlaxoSmithKline Intellectual Property Development Ltd., Sanofi US Services Inc., Maze Therapeutics Inc., Janssen Biotech Inc, Novartis AG, and Boehringer Ingelheim International GmbH. Following biobanks are acknowledged for delivering biobank samples to FinnGen: Auria Biobank (www.auria.fi/biopankki), THL Biobank (www.thl.fi/biobank), Helsinki Biobank (www.helsinginbiopankki.fi), Biobank Borealis of Northern Finland (https://www.ppshp.fi/Tutkimus-ja-opetus/Biopankki/Pages/Biobank-Borealis-briefly-in-English.aspx), Finnish Clinical Biobank Tampere (www.tays.fi/en-US/Research_and_development/Finnish_Clinical_Biobank_Tampere), Biobank of Eastern Finland (www.ita-suomenbiopankki.fi/en), Central Finland Biobank (www.ksshp.fi/fi-FI/Potilaalle/Biopankki), Finnish Red Cross Blood Service Biobank (www.veripalvelu.fi/verenluovutus/biopankkitoiminta), Terveystalo Biobank (www.terveystalo.com/fi/Yritystietoa/Terveystalo-Biopankki/Biopankki/) and Arctic Biobank (https://www.oulu.fi/en/university/faculties-and-units/faculty-medicine/northern-finland-birth-cohorts-and-arctic-biobank). All Finnish Biobanks are members of BBMRI.fi infrastructure (https://www.bbmri-eric.eu/national-nodes/finland/). Finnish Biobank Cooperative -FINBB (https://finbb.fi/) is the coordinator of BBMRI-ERIC operations in Finland. The Finnish biobank data can be accessed through the Fingenious® services (https://site.fingenious.fi/en/) managed by FINBB. We greatly thank all UK Biobank and FinnGen participants, as well as the principal investigators, laboratory personnel and data management teams behind these efforts. This work was finically supported by the University of Helsinki Doctoral Programme in Population Health (Y.F.), the Research Council of Finland (315589 and 320129 to T.T.; 338507, 336825, and 352795 to M.P.), the HiLIFE Fellows Program (T.T.), and Sigrid Jusélius Foundation (T.T. and M.P.). Figure S1 was created with BioRender (https://www.biorender.com/).

## Author contributions

Conceptualization: T.T., M.P., Y.F.; acquisition, analysis, and intepretation of data: T.T., M.P., Y.F., A.K., S.R.; drafting of the manuscript: Y.F. and T.T.; crital revision and editing of the manuscript: T.T., M.P., Y.F.; visualization: Y.F.; supervision: T.T. and M.P.

## Web resources

UKB SNP-Heritability Browser, https://nealelab.github.io/UKBB_ldsc/h2_browser.html

BOLT-LMM, https://alkesgroup.broadinstitute.org/BOLT-LMM/BOLT-LMM_manual.html

GENESIS, https://github.com/yandorazhang/GENESIS

LAVA paritioning algorithm, https://github.com/cadeleeuw/lava-partitioning

LD score regression, https://github.com/bulik/ldsc

linemodels, https://github.com/mjpirinen/linemodels

FINEMAP, http://www.christianbenner.com/

REGENIE v2.2.4 pipeline, https://github.com/FINNGEN/regenie-pipelines

PLINK 1.9, www.cog-genomics.org/plink/1.9/

1000 Genomes Project phase 3, https://ftp.1000genomes.ebi.ac.uk/vol1/ftp/release/20130502/

HapMap phase 3, ftp://ftp.ncbi.nlm.nih.gov/hapmap/phase_3

## Data and Code availability

Researchres can access individual-level data from the UKB (https://www.ukbiobank.ac.uk/enable-your-research/apply-for-access) and FinnGen (https://www.finngen.fi/en/researchers/accessing) following the corrsponding data application procedures. Work performed using UKB data was done under application 22627. The code used in this study is available at: https://github.com/yufugen/DC_GWAS.

