## Supplemental Document for "Role of X chromosome and dosage compensation mechanisms in complex trait genetics"

### Supplemental Figures

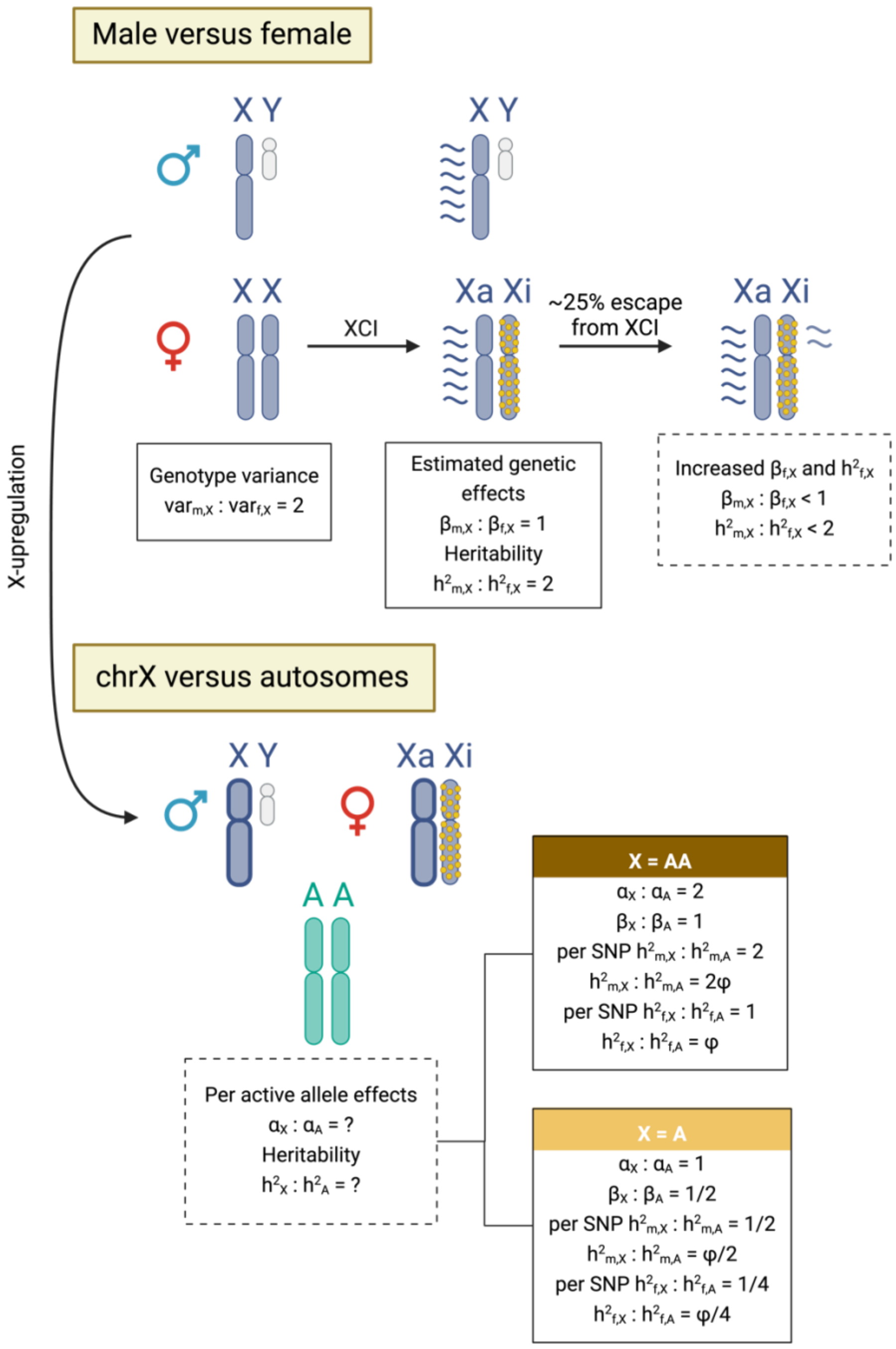


Figure S1. Illustration of the complications and consequences introduced by dosage compensation in the analysis and interpretation of genetic data in chrX. The assumptions are based on the model where the chrX alleles in females are coded as {0,1,2} and in males as {0,2}. $\beta_{f,X}$, $\beta_{m,X}$, female and male GWAS effects in chrX, respectively. $h^2_{f,X}$, $h^2_{m,X}$, chrX heritabilities in females and males, respectively. $a_X$, $a_A$, active allele effects in chrX and autosomes, respectively. $h^2_{f,A}$, $h^2_{m,A}$, autosomal heritabilities in females and males, respectively. $\phi$, the ratio of number of variants contributing to heritability in chrX versus that in autosomes. XCI, X chromosome inactivation.

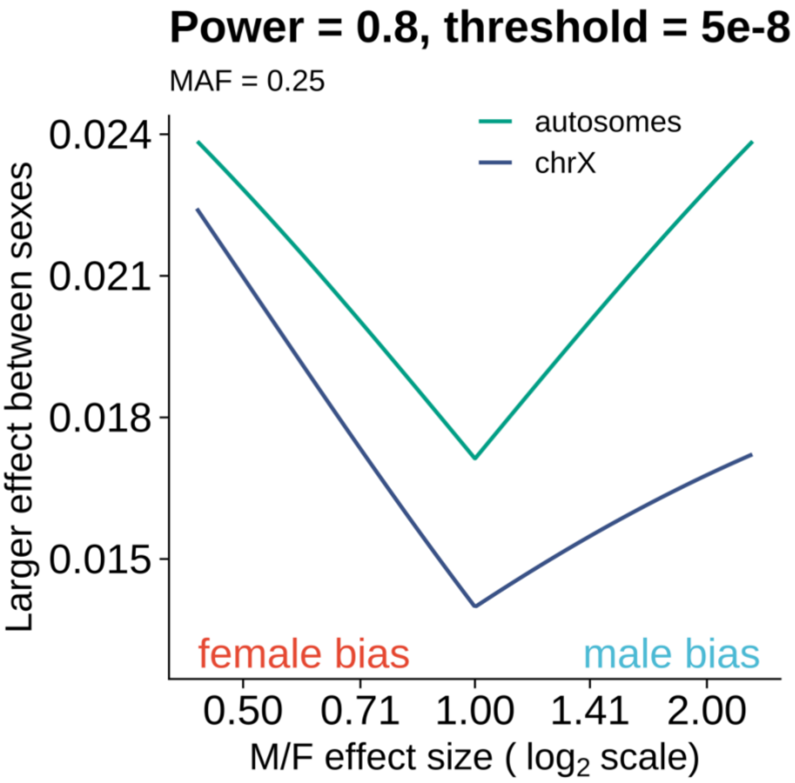


Figure S2. Illustration of the power bias in a sex-combined GWAS in chrX. How large the larger effect between sexes needs to be to be detected by a sex-combined GWAS ($n_m = n_f = 180{,}000$) with a power of 80% and a significance threshold at $5 \times 10^{-8}$ for a variant with minor allele frequency (MAF) at 0.25 with regards to different magnitude of sex difference in effect size.

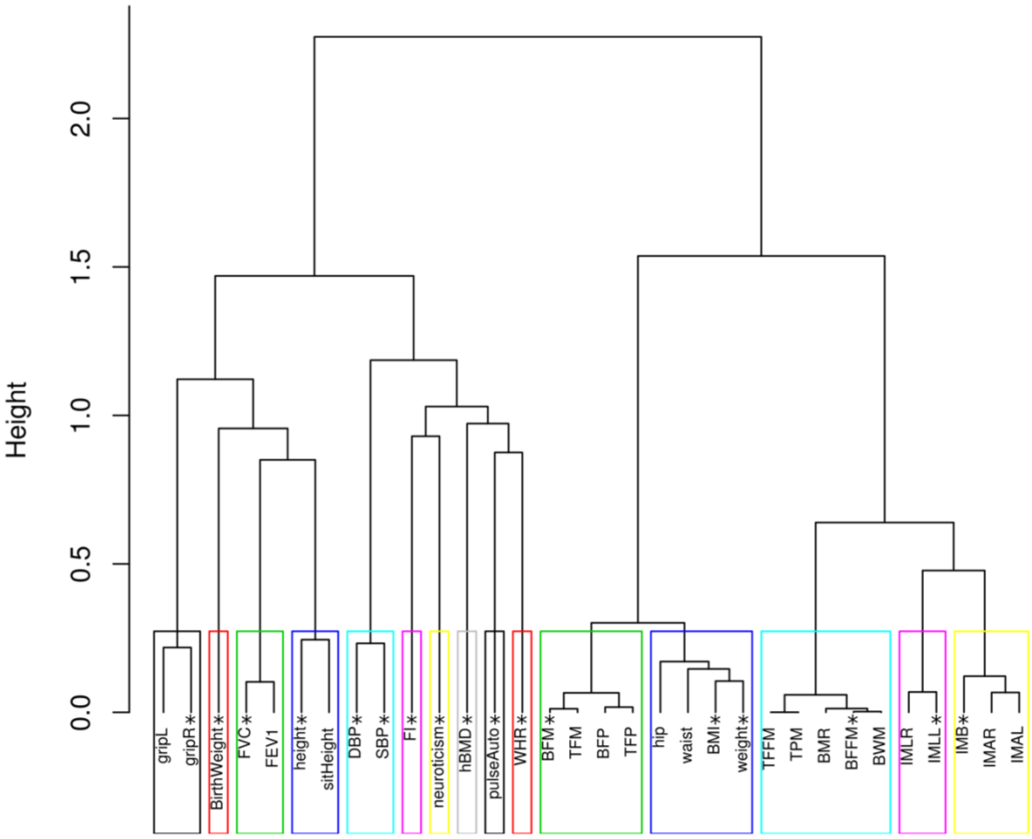


Figure S3. Hierarchical clustering dendrogram and identified clusters based on the correlation of adjusted and normalized trait values in the sex-combined population. The traits included in this study for GWAS are denoted with asterisks. Traits: hand grip strength, left (gripL) and right (gripR), birth weight (BirthWeight), forved vital capacity (FVC), forced expiratory volume in 1-second (FEV-1), standing height (height), sitting height (sitHeight), diastolic blood pressure (DBP), systolic blood pressure (SBP), fluid intelligence score (FI), neuroticism score (neuroticism), heel bone mineral density T-score (hBMD), automated reading pulse rate (pulseAuto), waist-to-hip ratio (WHR), whole body fat mass (BFM), trunk fat mass (TFM), body fat percentage (BFP), trunk fat percentage (TFP), hip circumference (hip), waist circumference (waist), body mass index (BMI), trunk fat-free mass (TFFM), trunk predicted mass (TPM), basal metabolic rate (BMR), whole body fat-free mass (BFFM), whole body water mass (BWM), impedance of leg, right (IMLR) and left (IMLL), impedance of whole body (IMB), impedance of arm, right (IMAR) and left (IMAL).

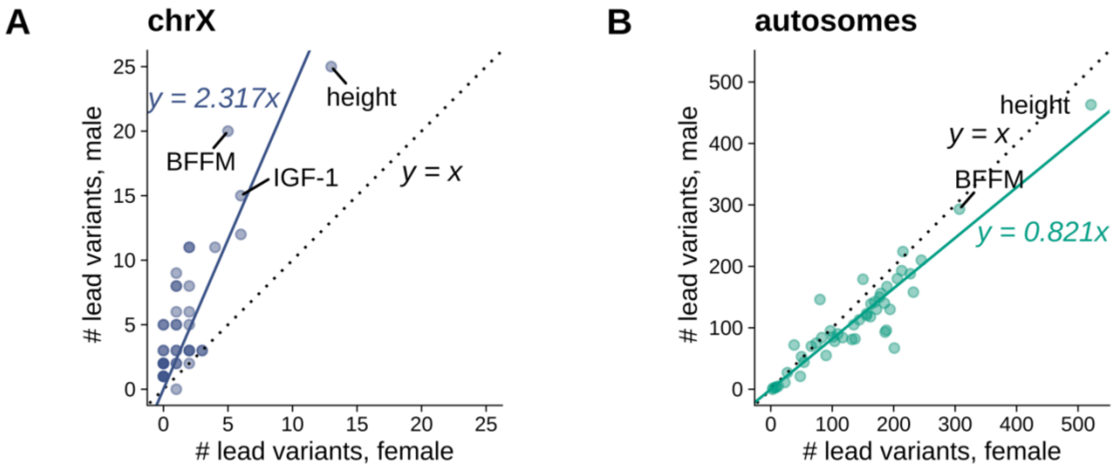


Figure S4. Comparison of the number of lead variants in male GWAS (y-axis) and female GWAS (x-axis) in (A) chrX and (B) autosomes across 48 traits. The dotted line indicates equal number of lead variants in male and female GWAS. The solid lines are regression lines. The numerical values are reported in Table S5. Abbreviations: whole body fat-free mass (BFFM), insulin-like growth factor 1 (IGF-1).

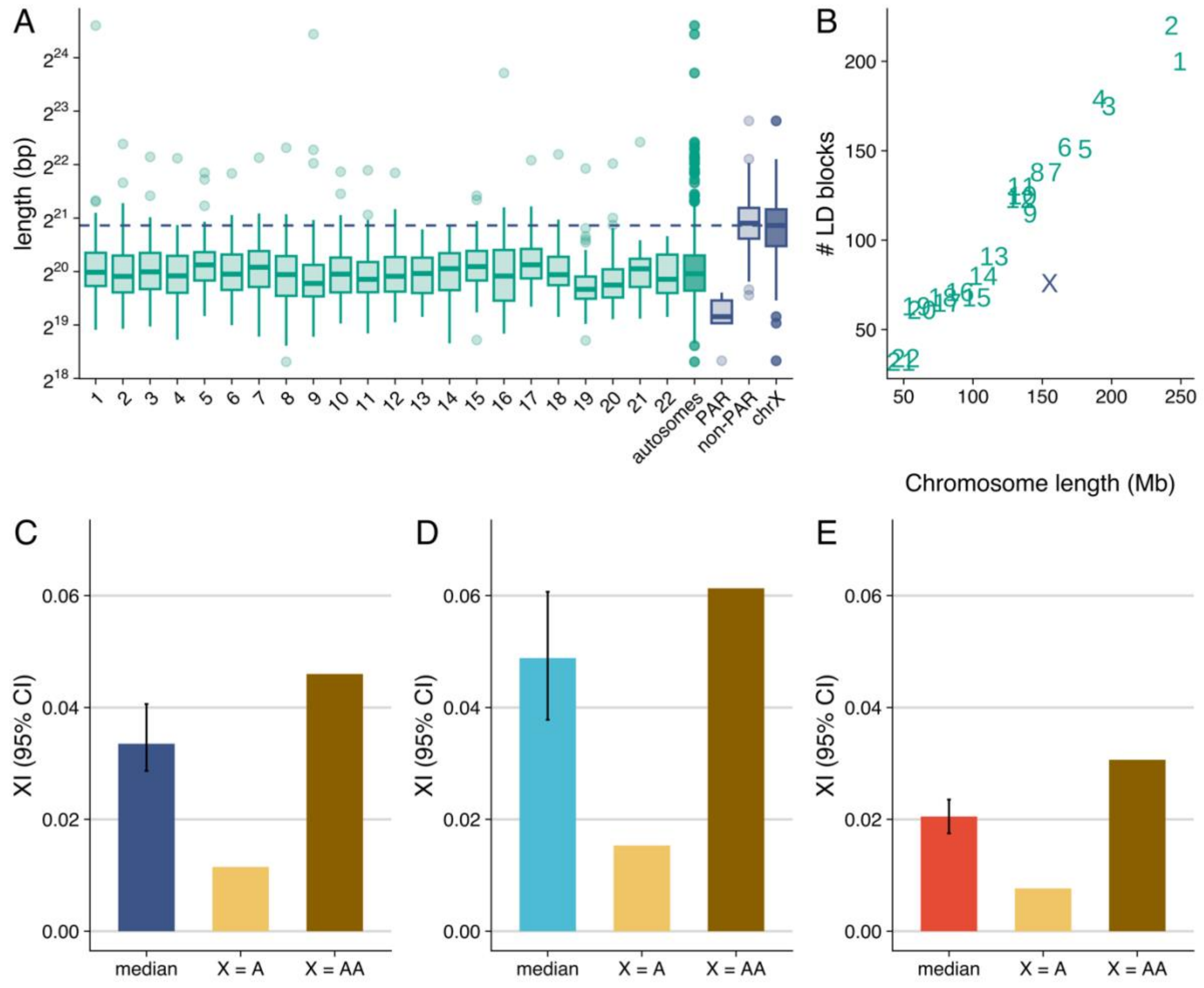


Figure S5. (A) Distribution of lengths of linkage disequilibrium (LD) blocks for each autosome, all autosomes, PAR and non-PAR of chrX and all chrX, shown as boxplots. The dashed line indicates the chrX median. The median LD block length in chrX is approximately twice that of the median autosome LD block length reflecting the reduced recombination in chrX. (B) The number of LD blocks in each chromosome versus the total length of the chromosome. Owing to the more extensive LD, the number of LD blocks in chrX is less than in autosomal chromosomes of similar length. (C-E) The median XI (bootstrap 95% CI) contrasted to the expected XI based on the number of LD blocks when the genetic effect of one active chrX is equal to one autosome (X = A) or equal to a pair of autosomes (X =

AA) (C) in the overall population, (D) in males, and (E) in females for 35 traits with nonzero $h_X^2$ in both sexes. Numerical values are reported in Table S2.

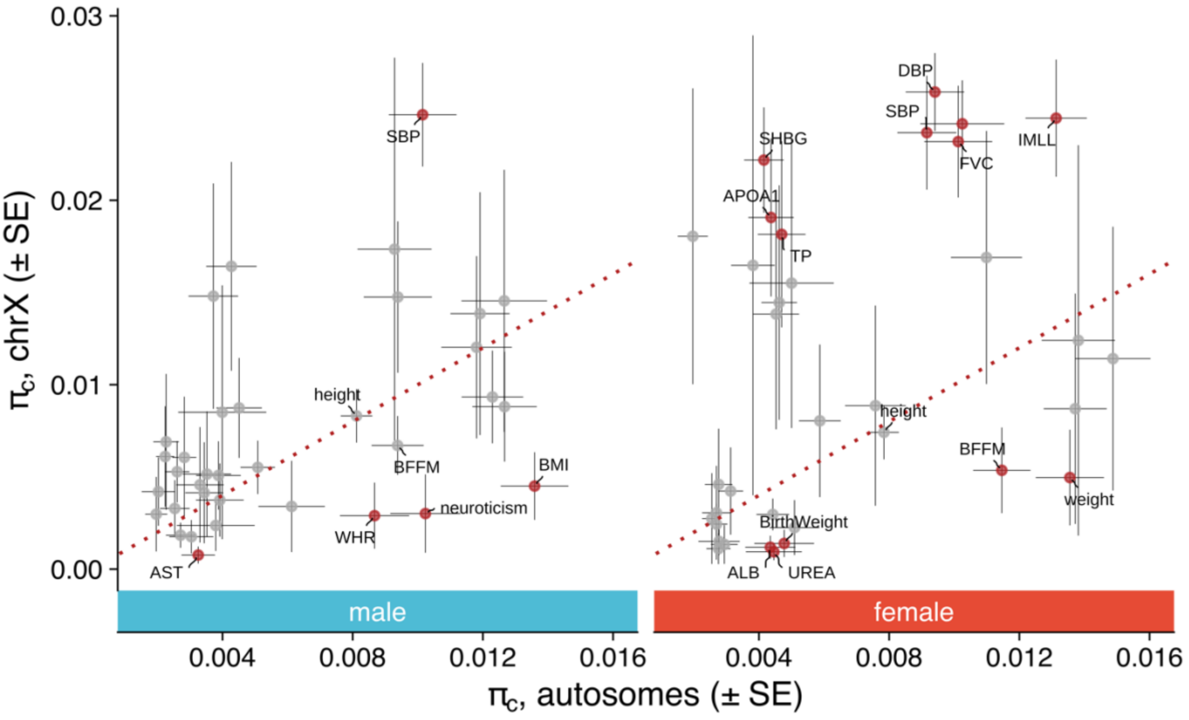


Figure S6. Comparison of the estimated proportion of causal variants ($\pi_c$) in chrX and in autosomes across 35 traits with nonzero $h_X^2$ in both sexes. Results are shown separately in males and in females. The dotted line indicates equal proportion of causal variants in chrX and autosomes. Numerical values reported in Tables S4 and S5. Abbreviations: systolic blood pressure (SBP), diastolic blood pressure (DBP), whole body fat-free mass (BFFM), body mass index (BMI), wasit-to-hip ratio (WHR), aspartate aminotransferase (AST), impedance of leg, left (IMLL), forced vital capacity (FVC), sex hormone-binding globulin (SHBG), apolipoprotein A (APOA1), total protein (TP), albumin (ALB), urea (UREA).

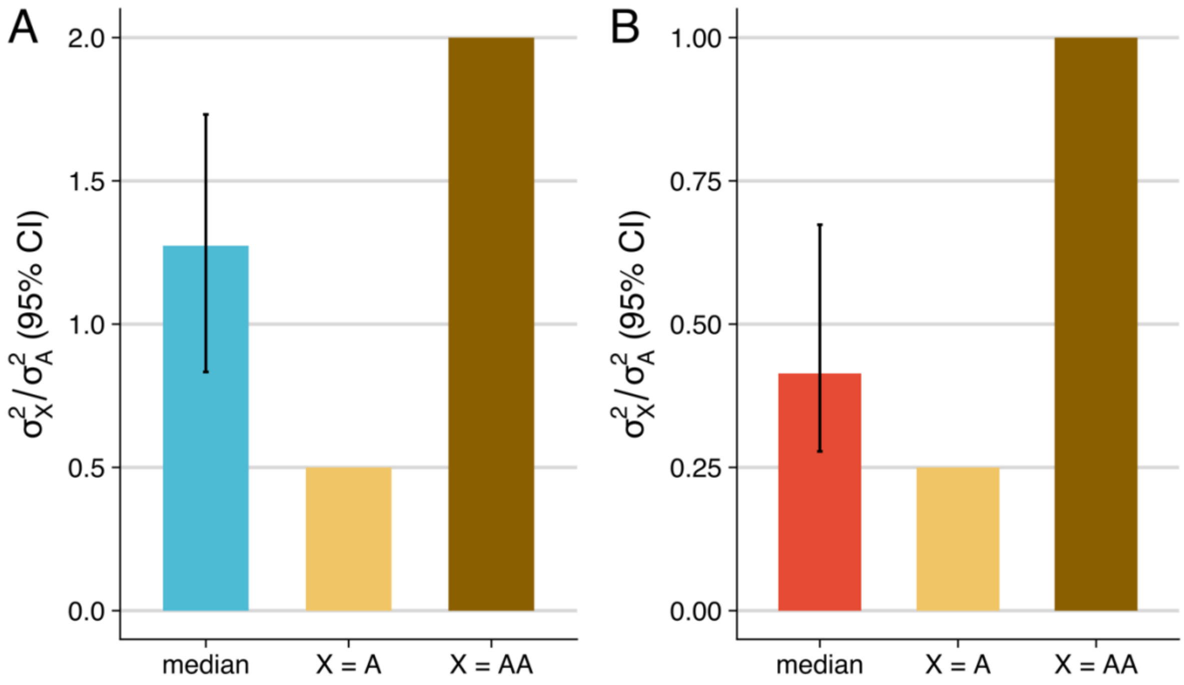


Figure S7. Comparison of $\sigma^2$ in chrX and in autosomes in (A) males and (B) females. The blue and red bars indicate the median of $\sigma_X^2/\sigma_A^2$ over the traits in males and females, respectively. The light and dark brown bars indicate the expected relationship between autosomal and chrX $\sigma^2$ under X = AA and X = A. Numerical values reported in Tables S4 and S5.

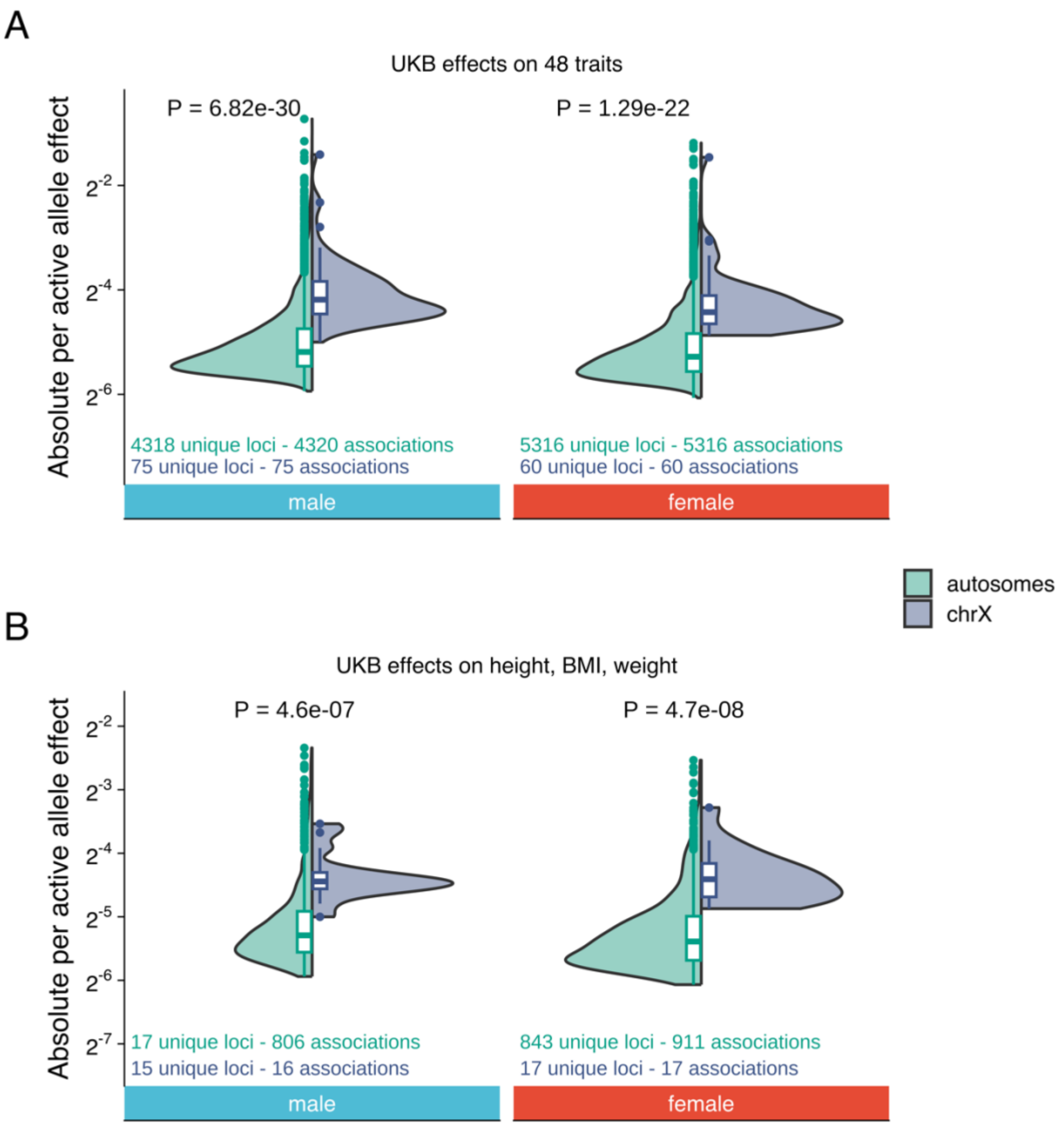


Figure S8. (A) Comparison of $a_X$ and $a_A$ for sex-specific unique trait-associated variants, where one trait was randomly selected for pleiotropic loci. (B) Comparison of $a_X$ and $a_A$ for variants associated with height, BMI, and weight in UKB with $a$ estimated in UKB (for comparison with Fig. 2B). Male GWAS in non-PAR have been down-sampled by half to achieve similar statistical power as in GWAS in autosomes. Numerical values are reported in Table S6 for UKB estimates and FinnGen summary statistics for FinnGen estimates.

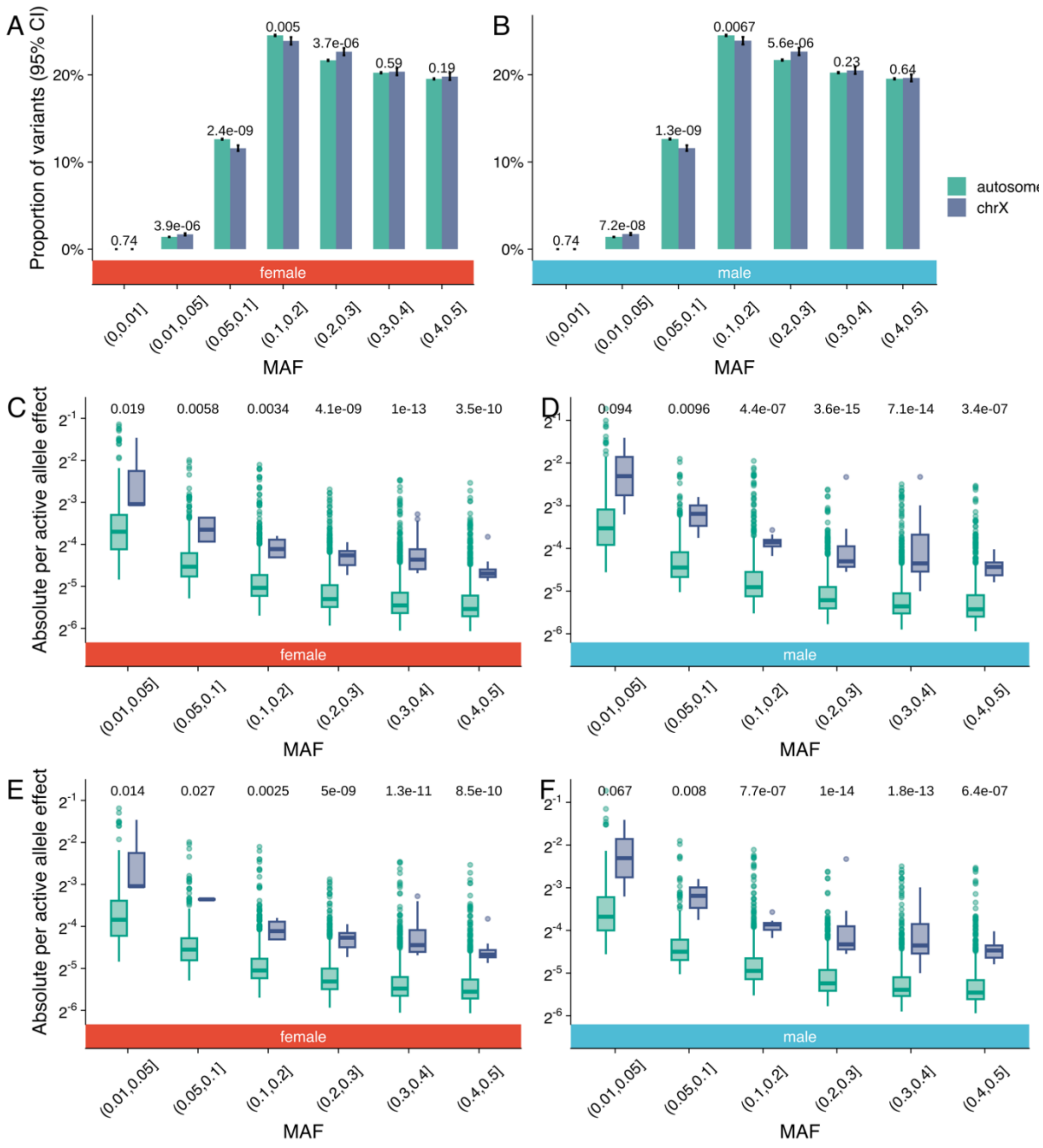


Figure S9. Comparison of MAF and the effects of MAF on $a$ between autosomes and chrX. We compared the MAF distributions of variants included in the GENESIS reference panel between autosomes and chrX, with MAF calculated in (A) female and (B) male UKB samples. *P*-values of proportion difference between autosomes and chrX ($\chi^2$ test) are indicated on top of each pair of bars. The distribution of $a$ of (C) female and (D) male lead variants for each MAF bin in autosomes and chrX across all associations. The male GWAS in non-PAR has been down-sampled by half. The distribution of $a$ of (E) female and (F) male lead variants for each MAF bin in autosomes and chrX, where one trait was randomly selected for pleiotropic variants. *P*-values of difference between $a_X$ and $a_A$ (Wilcox rank-sum test) were indicated on the top of the figures. Numerical values are reported in Table S6 for lead variants.

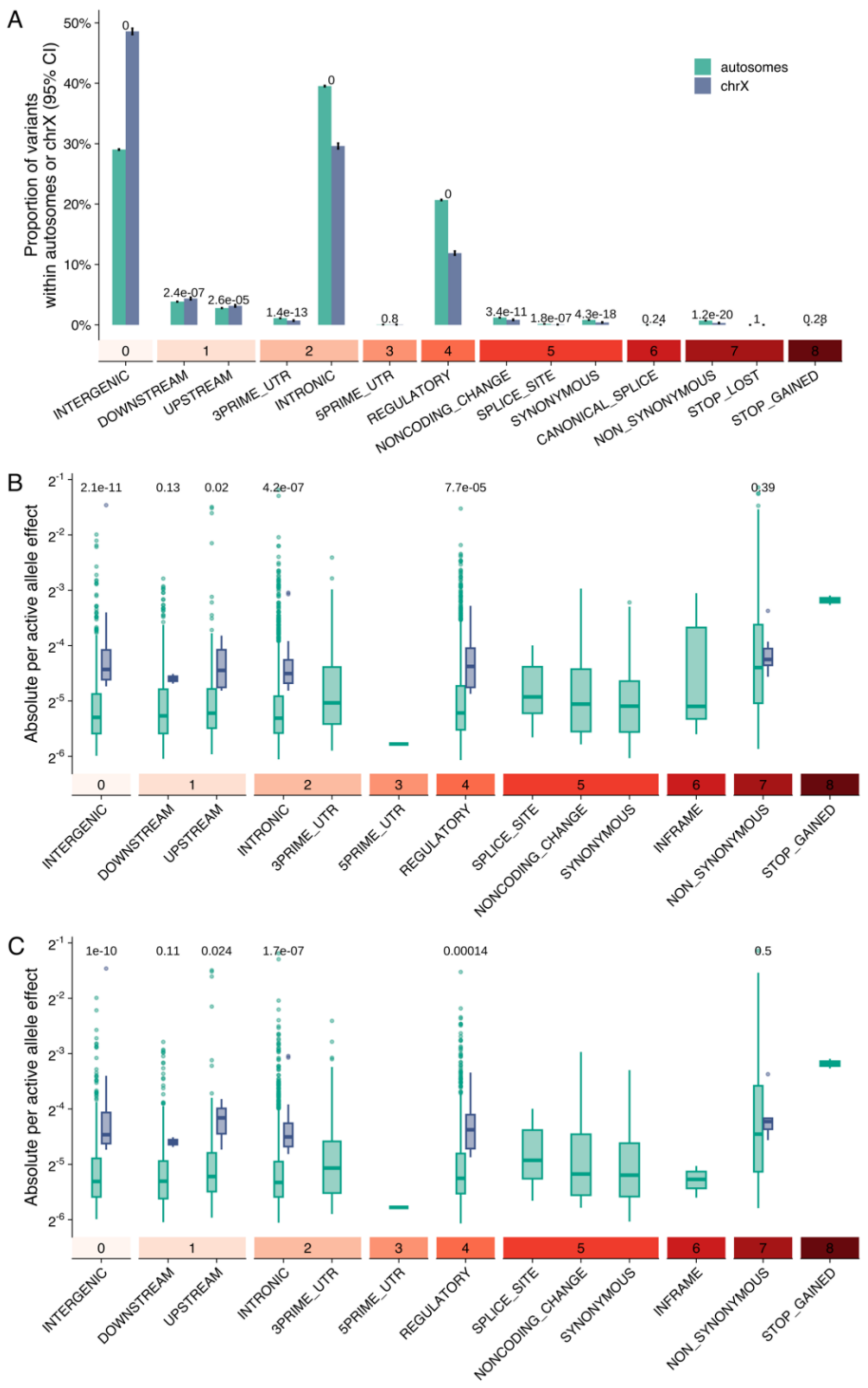


Figure S10. Comparison of functional consequences and the effects of functional consequences on active allele effect $a$ between autosomes and chrX. We compared (A) the proportions of functional consequences between common variants used in the GENESIS reference panel between autosomes and chrX. *P*-values of proportion difference between autosomes and chrX ($\chi^2$ test) are indicated on top of each pair of bars. We compared the $a_X$ and $a_A$ of female lead variants (B) across all associations and (C) with a single effect randomly selected for pleiotropic variants in the same functional consequence. *P*-values of difference between $a_X$ and $a_A$ (Wilcox rank-sum test) were indicated on the top of the figures. Numerical values are reported in Table S6 for lead variants. The numerical values and color of the blocks on top of the functional consequences indicate the severity of the consequences.

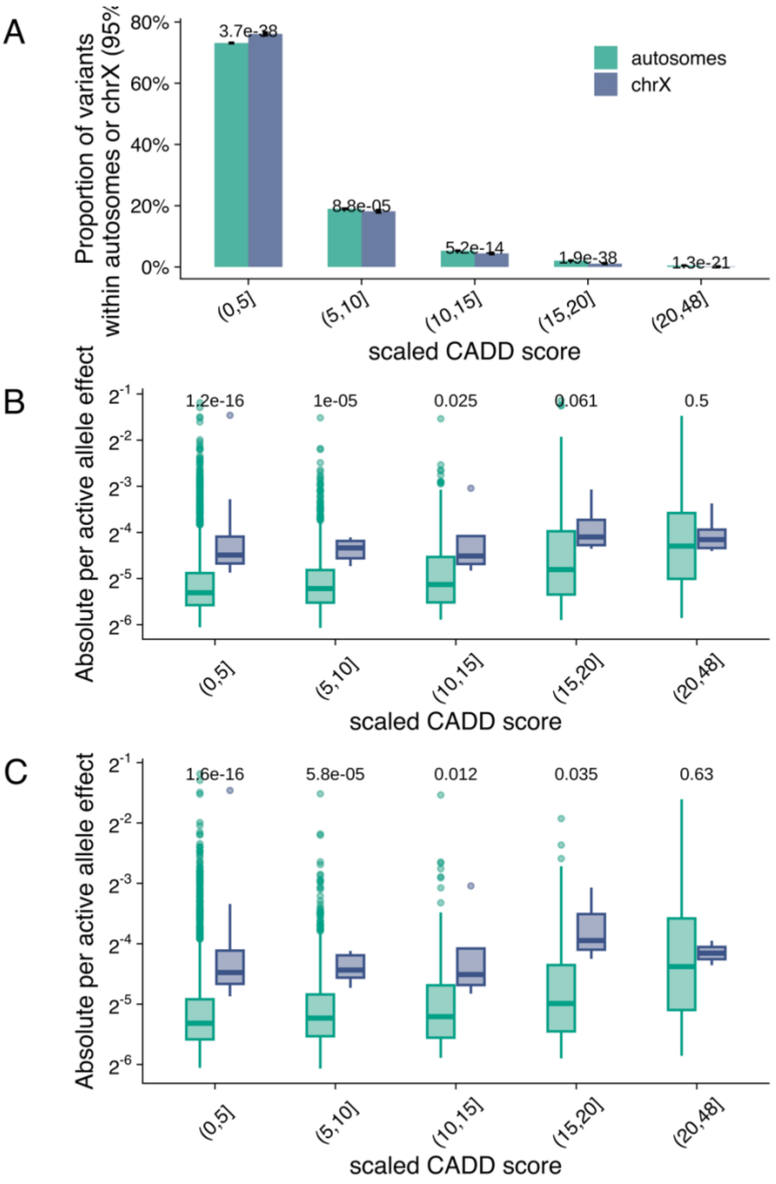


Figure S11. Comparison of pathogenicity measured as scaled CADD score and the effect of pathogenicity on $a$ between autosomes and chrX. The higher the scaled CADD score, the more pathogenic the variant is predicted to be. We compared (A) the proportions of scaled CADD scores between variants included in the GENESIS reference panel in autosomes and in chrX. *P*-values of proportion difference between autosomes and chrX ($\chi^2$ test) were indicated on top of each pair of bars. We compared (B) the $a_X$ and $a_A$ of female lead variants within in the same CADD bin and (C) with a single $a$ randomly selected for pleiotropic variants within each bin. *P*-

values of difference between $a_X$ and $a_A$ (Wilcox rank-sum test) were indicated on the top of the figures. Numerical values are reported in Table S6 for lead variants.

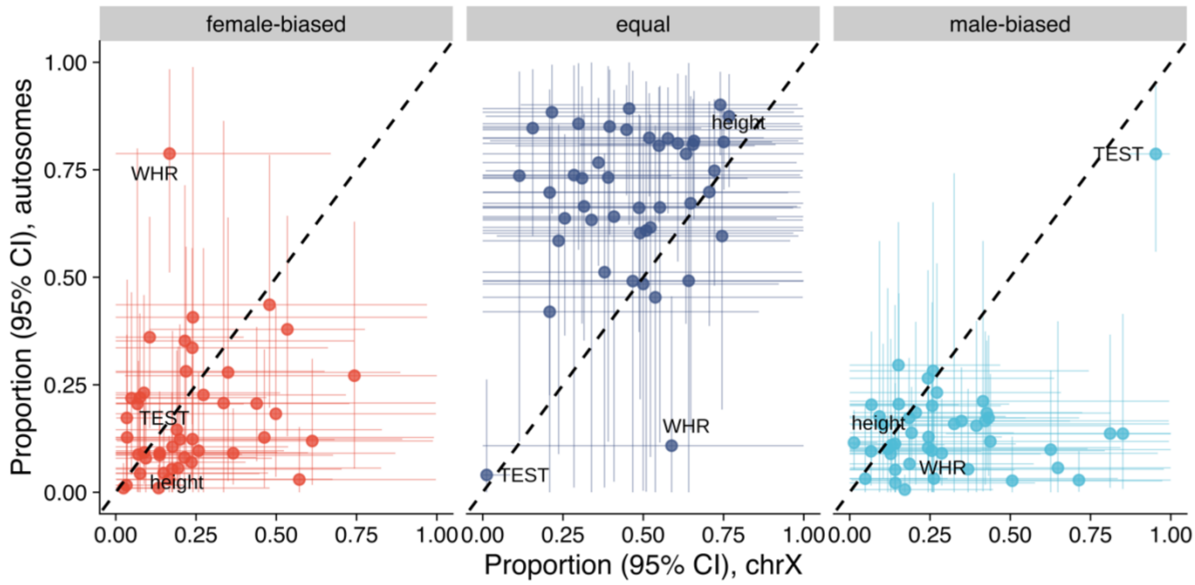


Figure S12. Comparison of the estimated non-null proportions of female-biased, equal, and male-biased components between chrX and autosomes. Numerical values are reported in Table S9. Abbreviations: waist-to-hip ratio (WHR), testosterone (TEST).

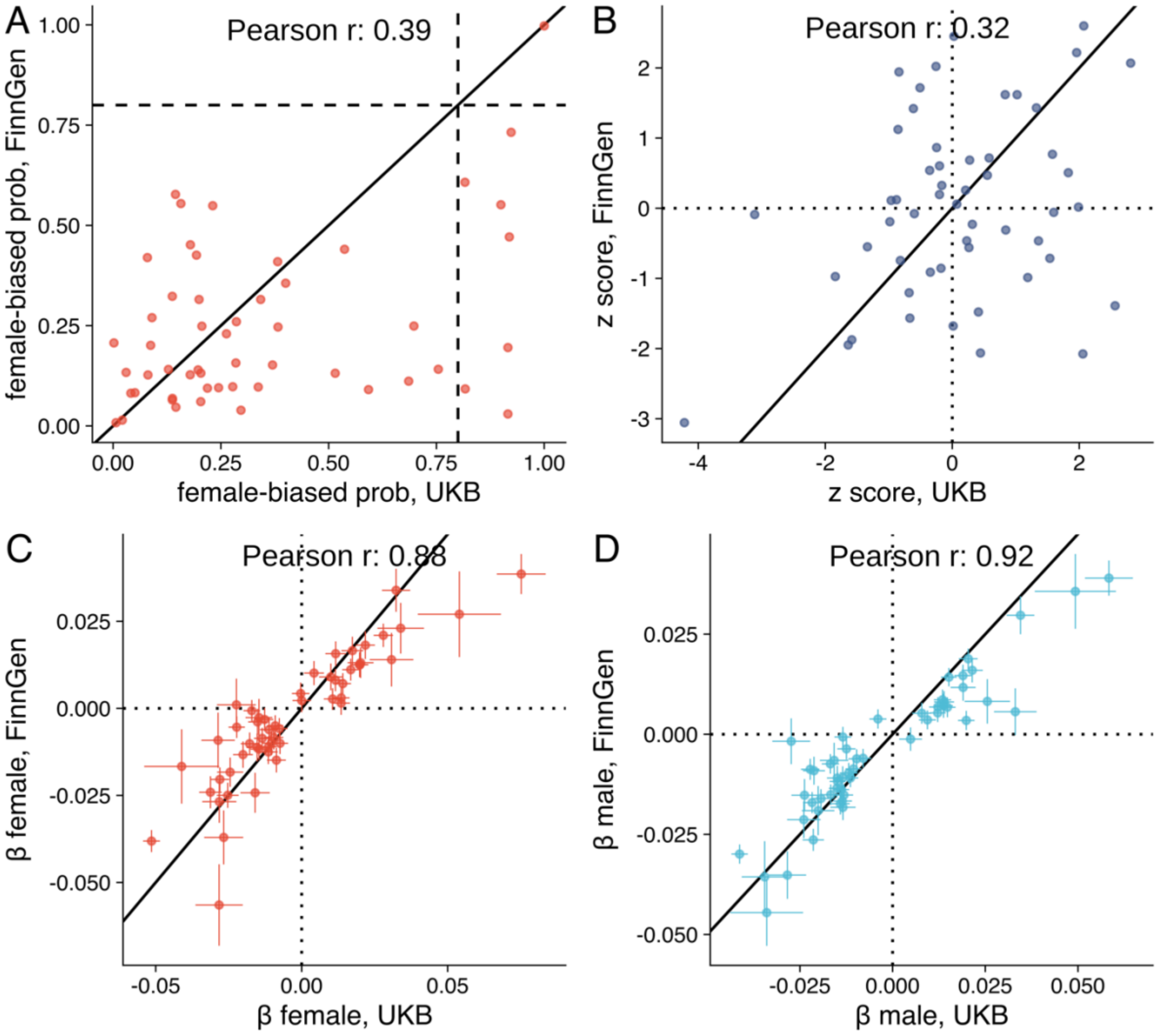


Figure S13. Comparison of UKB and FinnGen results in (A) female-biased probability, (B) sex difference z-score, (C) female effect sizes (± SE) and (D) male effect sizes (± SE) of lead variants identified in UKB height sex-combined GWAS in chrX. Numerical values are reported in Tables S12 and S13.

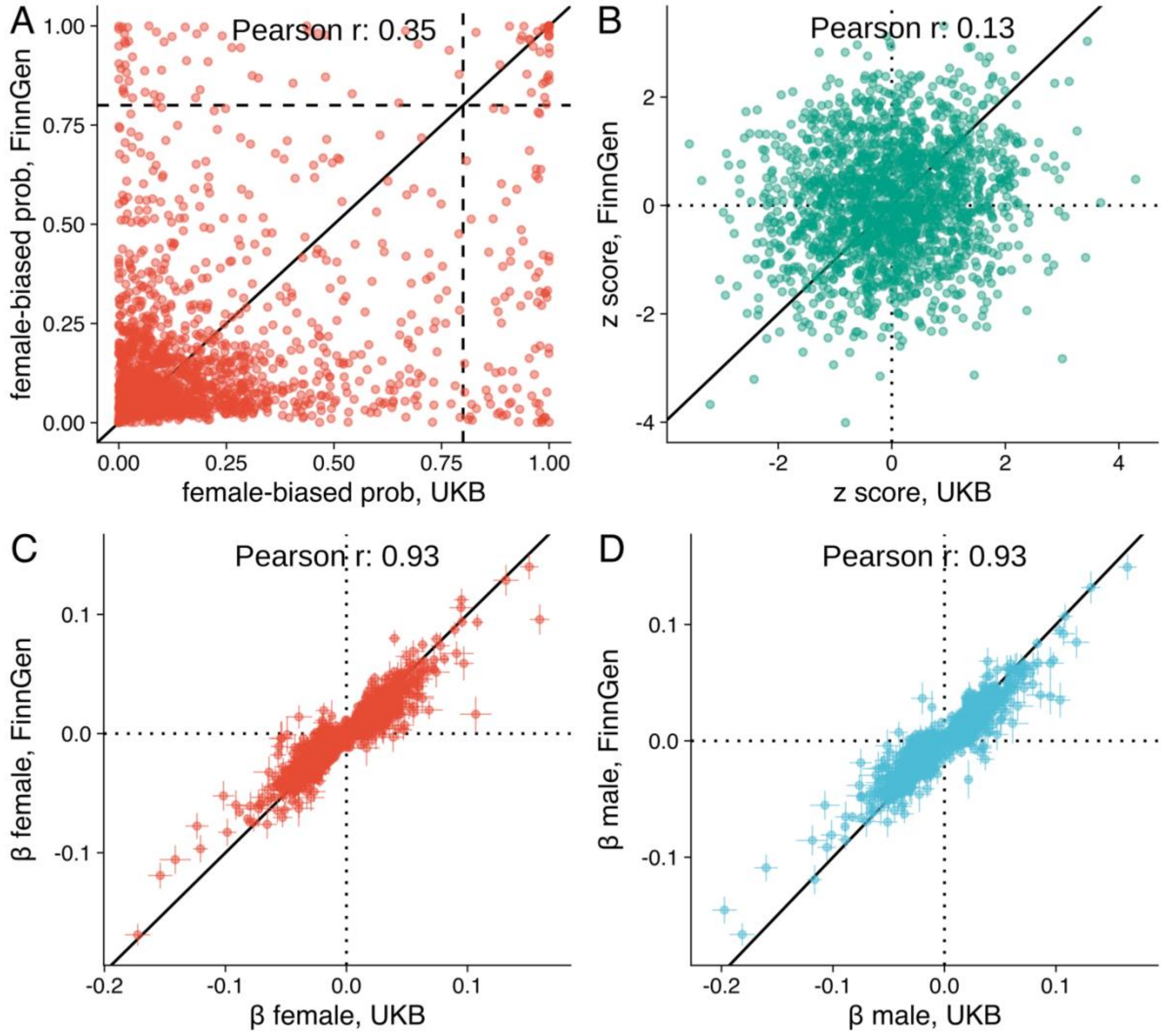


Figure S14. Comparison of UKB and FinnGen results in (A) female-biased probability, (B) sex difference z-score, (C) female effects (± SE) and (D) male effects (± SE) of lead variants identified in UKB height sex-combined GWAS in autosomes. Numerical values are reported in Tables S12 and S13.

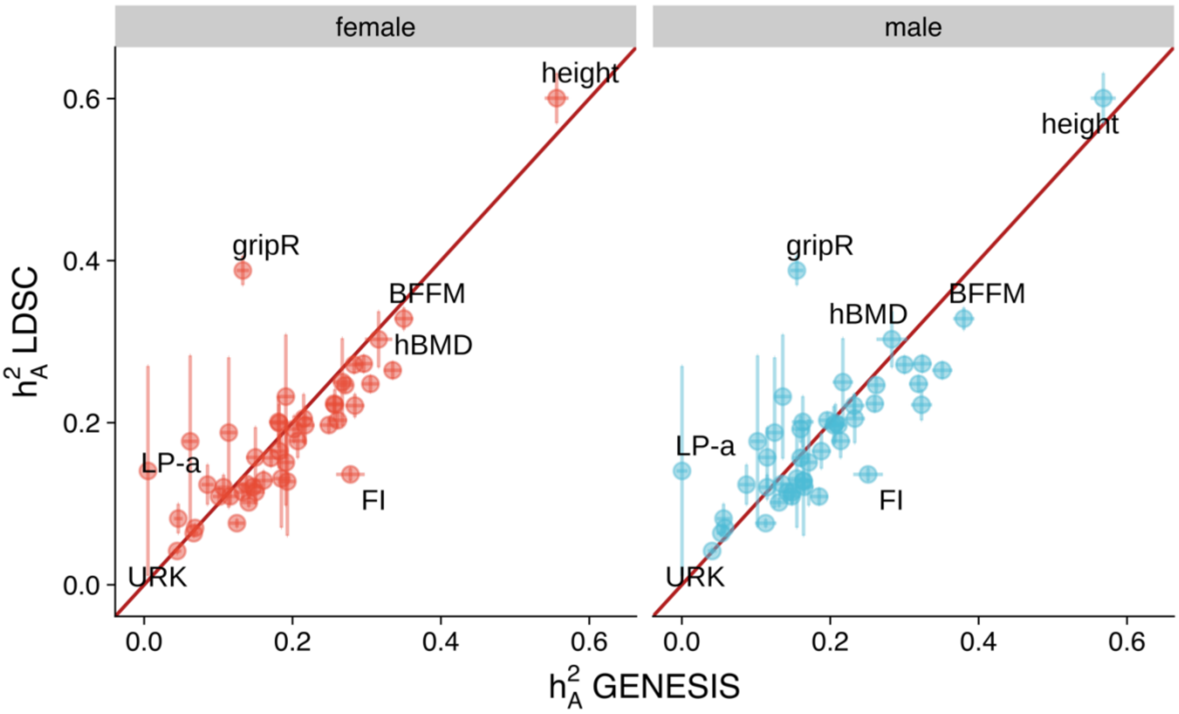


Figure S15. Comparison of estimates of autosomal heritability $h_A^2$ for 48 traits. We compared the estimated $h_A^2$ by GENESIS to that estimated by LDSC using female and male summary statistics. Error bars indicate SE of $h_A^2$. Solid lines indicate equal estimates between the two methods. Numerical results are in Table S2. Abbreviations: hand grip strength, right (gripR), whole body fat-free mass (BFFM), heel bone mineral density T-score (hBMD), lipoprotein A (LP-a), fluid intelligence score (FI), potassium in urine (URK).

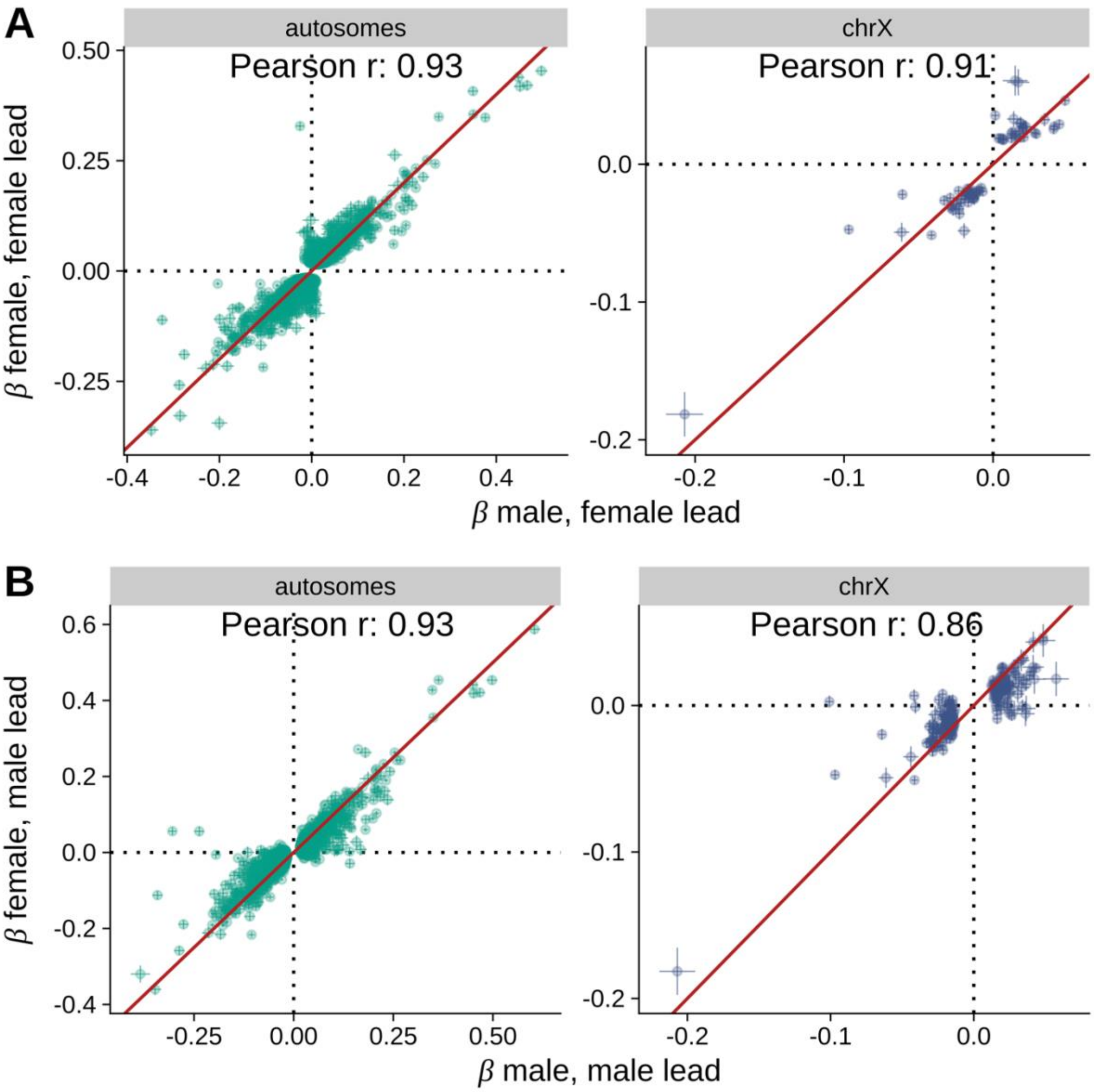


Figure S16. The male and female effects (± SE) of lead variants from (A) female-specific GWAS and (B) male-specific GWAS. The Pearson correlations after removing variants associated with testosterone, a trait known for its sex-specific effects and strong contribution from chrX, were 0.94 in autosomes and 0.92 in chrX in females and 0.96 in autosomes and 0.91 in chrX in males. The numerical values are reported in Table S6.

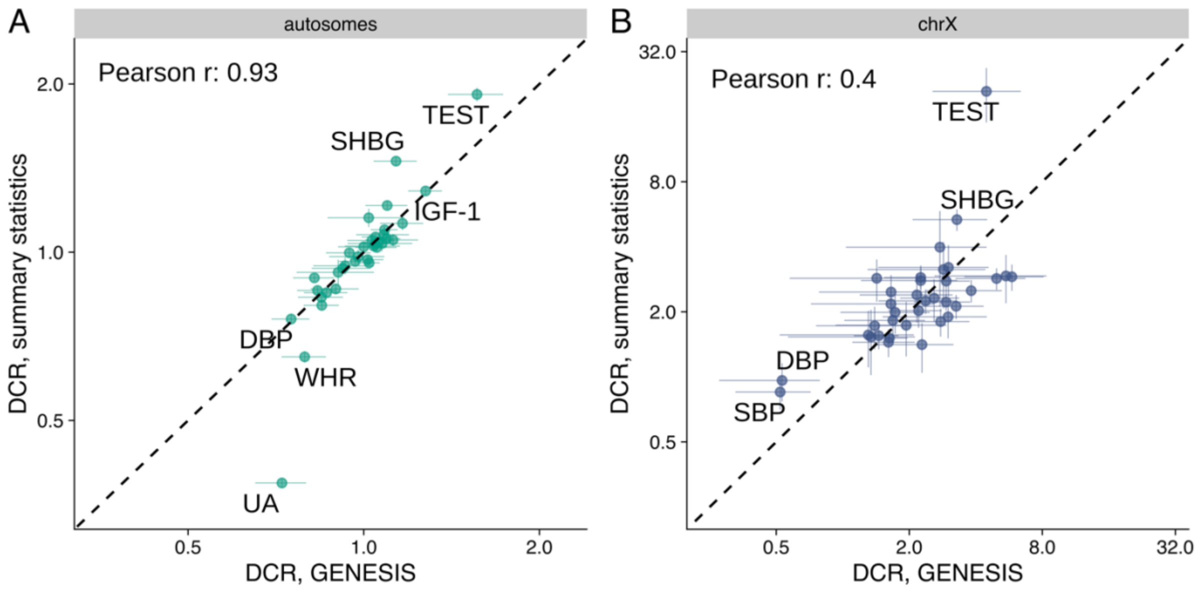


Figure S17. Comparison of autosomal and chrX dosage compensation ratio (DCR) using different methods. We compared DCR (± SE) estimated using summary statistics and using $h^2$ estimated from GENESIS for (A) autosomes and (B) chrX. Numerical values are reported in Table S2. Abbreviations: testosterone (TEST), sex hormone-binding globulin (SHBG), insulin-like growth factor 1 (IGF-1), diastolic blood pressure (DBP), systolic blood pressure (SBP), waist-to-hip ratio (WHR), urate (UA).

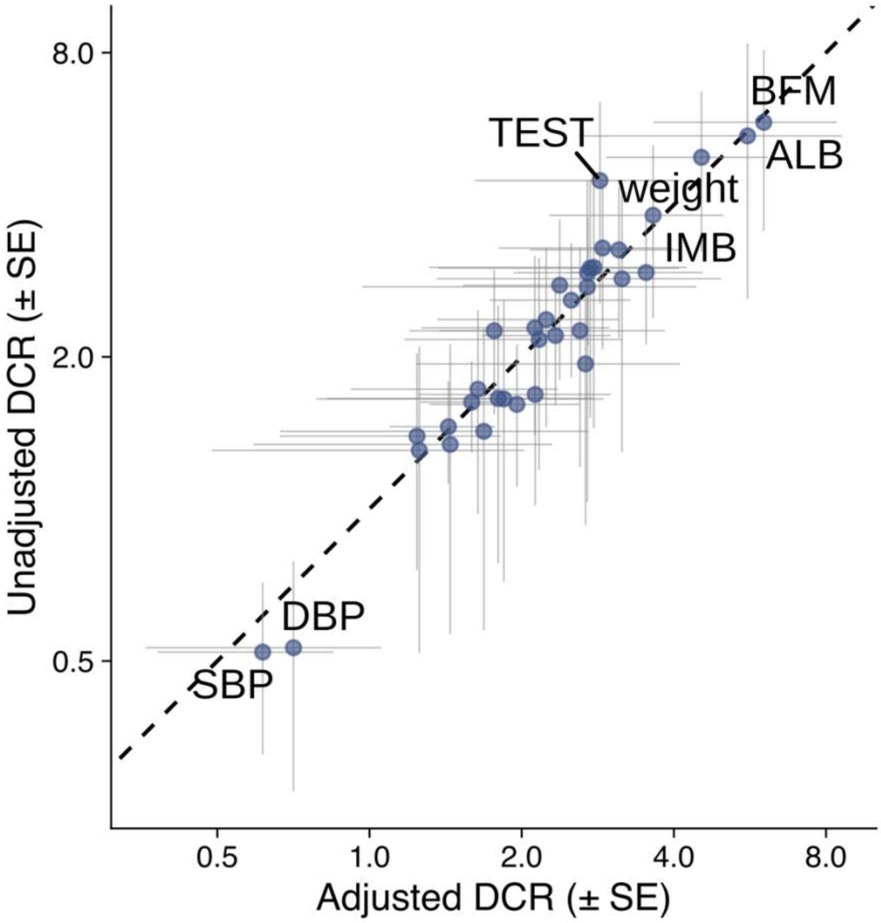


Figure S18. Comparison of chrX dosage compensation ratio (DCR) estimates with and without adjustment for autosomal DCR estimates. Numerical values are reported in Table S2. Abbreviations: whole body fat mass (BFM), albumin (ALB), testosterone (TEST), impedance of whole body (IMB), diastolic blood pressure (DBP), systolic blood pressure (SBP).

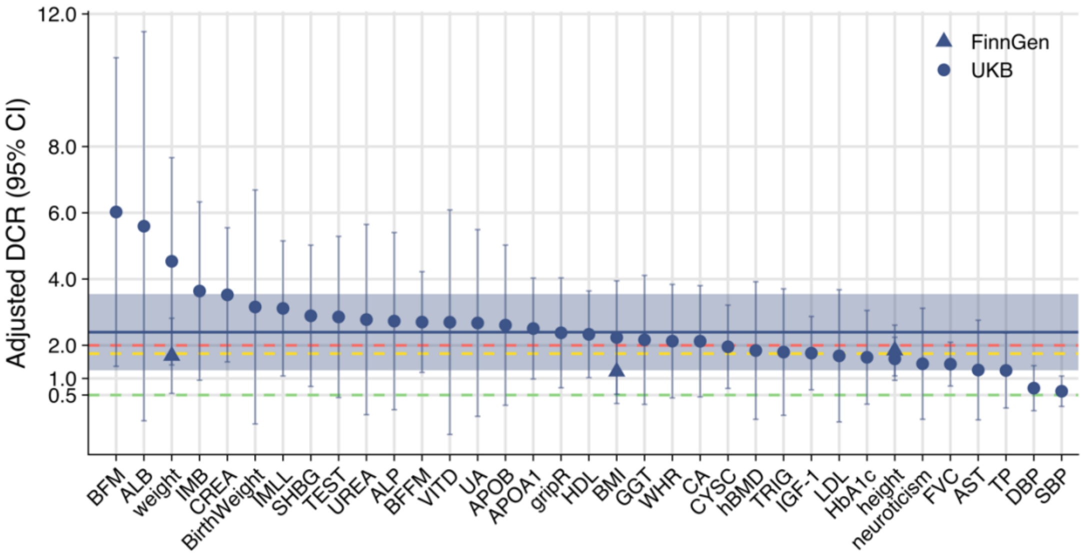


Figure S19. (A) ChrX dosage compensation ratio (DCR) estimates adjusted by autosomal DCR estimates with 95% CI (DCR ± 1.96*SE) for 35 traits with nonzero $h_X^2$ in both sexes using $h_X^2$ estimated by GENESIS. The solid blue line indicates the mean DCR and the shaded region indicates one standard deviation of the DCR point estimates of the traits. The red, yellow and green dashed lines indicate expectation under full XCI, 25% escape from XCI, and no XCI, respectively. Numerical values are reported in Table S2. Abbreviations: whole body fat mass (BFM), albumin (ALB), impedance of body (IMB), creatinine (CREA), impedance of leg, left (IMLL), sex hormone-binding globulin (SHBG), testosterone (TEST), urea (UREA), alkaline phosphatase (ALP), whole body fat-free mass (BFFM), vitamin D (VITD), urate (UA), apolipoprotein B (APOB), apolipoprotein (APOA1), hand grip strength, right (gripR), high-density lipoprotein cholesterol (HDL), body mass index (BMI), gamma glutamyltransferase (GGT), waist-to-hip ratio (WHR), calcium (CA), cystatin C (CYSC), heel bone mineral density T-score (hBMD), triglycerides (TRIG), insulin-like growth factor 1 (IGF-1), low-density lipoprotein cholesterol (LDL), glycated haemoglobin (HbA1c), forced vital capacity (FVC), aspartate aminotransferase

(AST), total protein (TP), diastolic blood pressure (DBP), systolic blood pressure (SBP).

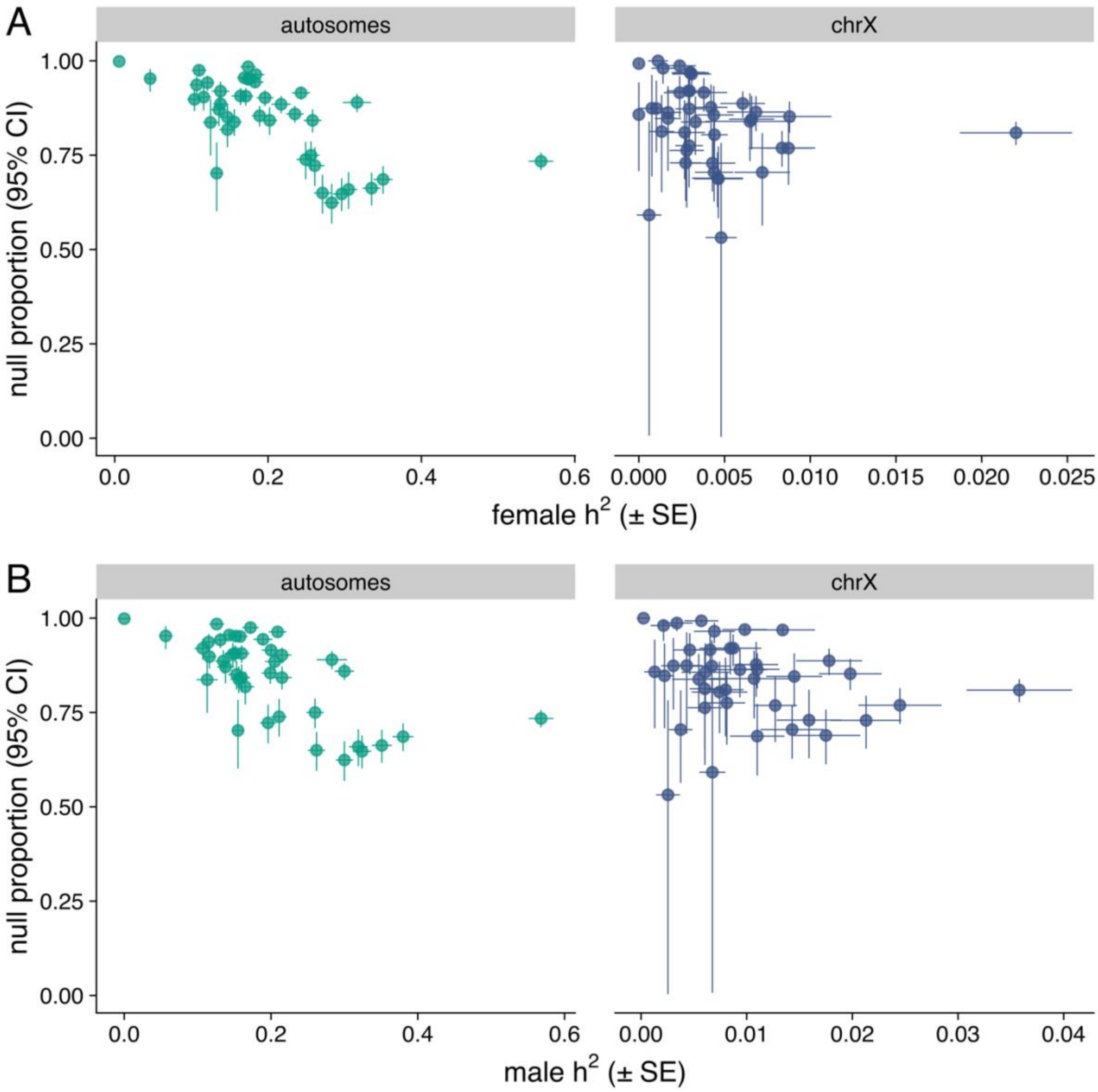


Figure S20. Comparison of estimated proportion of null variants using the mixture model of sex-specific effect sizes for female (A) and male (B) $h^2$ estimated with GENESIS. Numerical values are reported in Tables S2 and S9.

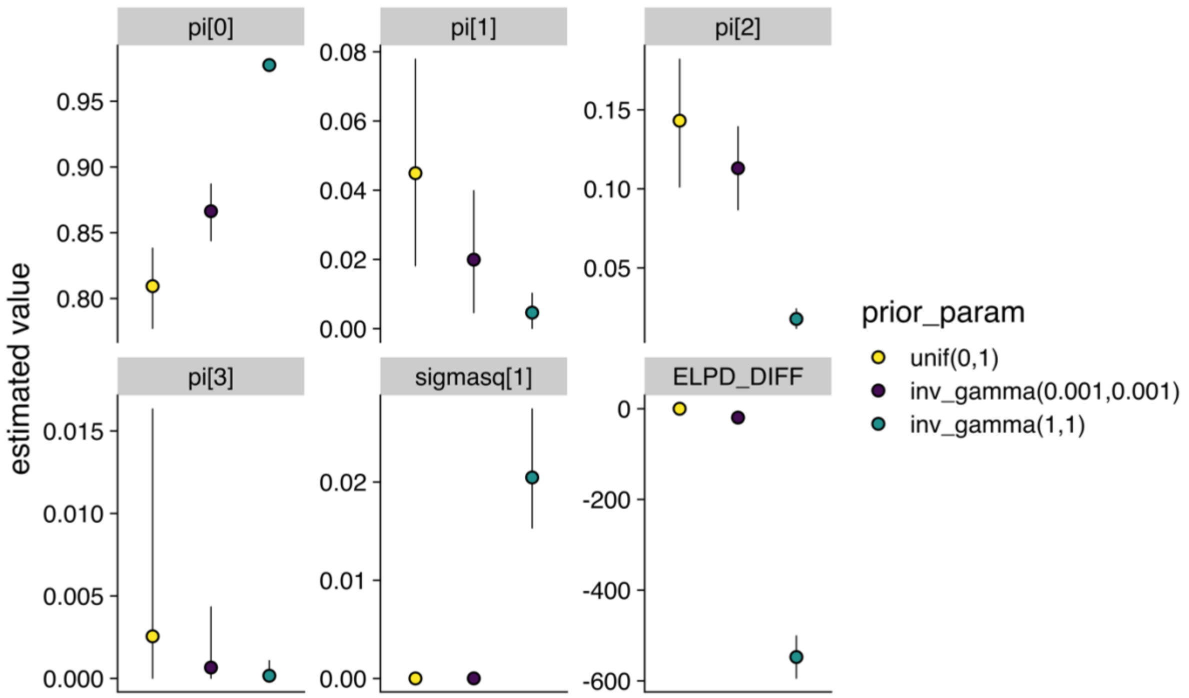


Figure S21. Comparison of different priors on $\sigma^2$ in four-component mixture model to classify variants between null, equal, female-biased and male-biased components. Estimated posterior distributions of 5 parameters and ELPD-DIFF values with 3 different priors for $\sigma^2$. For estimated proportion parameters $\pi_1, \ldots, \pi_4$ and non-zero effects' variance $\sigma^2$ means and 2.5 to 97.5 percentile intervals are shown. For ELPD-DIFF, which is the difference in ELPD-LOO of each model relative to the model with the highest ELPD-LOO, is shown together with its SE. Here, unif(0,1) prior has the highest ELPD-LOO and is therefore preferred. Numeric values are reported in Tables S10 and S11. See https://mc-stan.org/loo/reference/loo_compare for a description of this model assessment approach.

#### Supplemental Tables

Table S1. Information on the UKB and FinnGen traits and the covariates used in genome-wide association analysis (GWAS).

Table S2. Estimated sex-specific and sex-combined $h^2$ in autosomes and chrX with GENESIS and LDSC. Sex-combined estimate is computed as average of sex-specific estimates. We tested if $h^2$ is different from zero (indicated by the “h2_x_pval” column) using GENESIS estimates and also report FDR adjusted *P*-values. "XI" is the chrX-to-autosome $h^2$ ratio, with "XI_se" the corresponding standard error (SE). We compared the male and female $h^2$ estimates in chrX and autosomes, with “pval_sex_diff_x” and “pval_sex_diff_a” reporting the corresponding *P*-values in chrX and autosomes, and “fdr_sex_diff_x” and “fdr_sex_diff_a” the corresponding FDR adjusted *P*-values. “DCR_GENESIS” is calculated based on GENESIS estimates for chrX and autosomes. “DCR_sumstats” is calculated using summary statistics.

Table S3. The effect size distribution estimates from GENESIS M2 model in autosomes and chrX with female summary statistics. The proportions of causal SNPs are in columns “pic.autosomes” and “pic.se.autosomes” for autosomes, and “pic.x” and “pic.se.x” for chrX. The per SNP $h^2$ are in columns “sigmasq.autosomes” and “sigmasq.se.autosomes” for autosomes, and “sigmasq.x” and “sigmasq.se.x” for chrX. The estimated $h^2$are in “h2.autosomes” and “h2.se.autosomes” for autosomes and “h2.x” and “h2.se.x” for chrX, which are the same as the estimated female $h^2$ by GENESIS in Table S2. The estimated number of causal SNPs are in columns “nbr.sSNP.autosomes” and “nbr.sSNP.se.autosomes” for autosomes and “nbr.sSNP.x” and “nbr.sSNP.se.x” for chrX.

Table S4. The effect size distribution estimates from GENESIS M2 model in autosomes and chrX with male summary statistics. The proportions of causal SNPs are in columns “pic.autosomes” and “pic.se.autosomes” for autosomes, and “pic.x” and “pic.se.x” for chrX. The per SNP $h^2$ are in columns “sigmasq.autosomes” and “sigmasq.se.autosomes” for autosomes, and “sigmasq.x” and “sigmasq.se.x” for chrX. The estimated $h^2$are in “h2.autosomes” and “h2.se.autosomes” for autosomes and “h2.x” and “h2.se.x” for chrX, which are the same as the estimated male $h^2$ by GENESIS in Table S2. The estimated number of causal SNPs are in columns “nbr.sSNP.autosomes” and “nbr.sSNP.se.autosomes” for autosomes and “nbr.sSNP.x” and “nbr.sSNP.se.x” for chrX.

Table S5. The number of sex-specific LD-independent lead variants (*P*-value < $5 \times 10^{-8}$) in autosomes and chrX.

Table S6. Summary statistics of lead variants identified in sex-specific GWAS for all traits. In the allelic effect columns “ALLELE_EFF_FEMALE” and “ALLELE_EFF_MALE” and corresponding SE in columns “ALLELE_SE_FEMALE” and “ALLELE_SE_MALE” the GWAS effect estimates “BETA_FEMALE” and “BETA_MALE”, “SE_FEMALE” and “SE_MALE” were multiplied by two for variants in the non-PAR region to correspond the active allele effects described in the main text. The samples included in each GWAS were indicated in the column “SAMPLE”, which is “female” for female-specific GWAS, “male” for full sample sized male-specific GWAS, “downsize male” for downsized male-specific GWAS in the non-PAR region. “ALLELE1” was used as the effect allele and “ALLELE0” the reference allele in GWAS. “A1FREQ_FEMALE” and “A1FREA_MALE” are the frequencies of the effective allele in females and males, respectively. And “MAF_FEMALE” AND “MAF_MALE” are the minor allele frequencies in females and males, respectively.

“P_BOLT_LMM_INF_FEMALE” and “P_BOLT_LMM_INF_MALE” are the BOLT-LMM infinitesimal mixed model association test *P*-values in females and males. “Consequence” is the VEP predicted consequence of the SNP and corresponding severity of the consequence “ConsScore” was reported by CADD, and the affected gene were given with Stable ID (“GeneID”) and gene name (“GeneName”). The CADD prediction of the pathogenicity of each variant is in columns “RawScore” for unscaled C-score and “PHRED” for scaled C-score.

Table S7. Estimated number of LD blocks and average length of the LD blocks per chromosome and aggregated over all autosomes.

| Chromosome | Chromosome length (bp) | Number of LD blocks | Average length of LD blocks (bp) |
|---|---|---|---|
| 1 | 249250621 | 200 | 1246150.0 |
| 2 | 243199373 | 220 | 1105355.4 |
| 3 | 198022430 | 175 | 1130869.6 |
| 4 | 191154276 | 179 | 1067227.1 |
| 5 | 180915260 | 151 | 1197965.9 |
| 6 | 171115067 | 168 | 1017794.2 |
| 7 | 159138663 | 138 | 1152996.4 |
| 8 | 146364022 | 138 | 1060087.3 |
| 9 | 141213431 | 115 | 1227271.4 |
| 10 | 135534747 | 125 | 1083710.9 |
| 11 | 135006516 | 130 | 1037379.2 |
| 12 | 133851895 | 123 | 1087654.5 |
| 13 | 115169878 | 91 | 1055931.4 |
| 14 | 107349540 | 80 | 1103617.2 |
| 15 | 102531392 | 68 | 1213540.8 |
| 16 | 90354753 | 71 | 1270882.0 |
| 17 | 81195210 | 65 | 1249114.0 |
| 18 | 78077248 | 68 | 1147152.3 |
| 19 | 59128983 | 63 | 937366.8 |
| 20 | 63025520 | 61 | 1031194.0 |
| 21 | 48129895 | 32 | 1209637.2 |
| 22 | 51304566 | 34 | 1035118.4 |
| Autosomes | 2985940020 | 2495 | 1120667.0 |
| chrX, non-PAR | 152301523 | 71 | 2144094.0 |
| chrX, PAR | 2969037 | 5 | 593267.0 |
| chrX | 155270560 | 76 | 2042066.0 |

Table S8. Posterior probabilities of XCI scenarios for each trait based on sex-specific $h^2$ in chrX and in autosomes.

Table S9. Estimated parameters, proportions of null (p[0]), female-biased (p[1]), equal (p[2]) and male-biased (p3) components and $\sigma^2$, of all traits in four-component sex bias mixture model. We report the summaries of the parameters: mean (“mean”), Monte Carlo standard errors (“se_mean”), standard deviations (“sd”), quantiles (“2.5%”,”50%” and “97.5%”), effective sample sizes (“n_eff”), split Rhats (“Rhat”). The “Data” column indicates whether UKB or FinnGen summary statistics were used.

Table S10. The estimates (mean), and 2.5, 50, 97.5 percentile (“2.5%”,”50%” and “97.5%”) of each parameter with different priors for sigmasq (“prior_param”) using UKB height summary statistics in chrX, as well effective sample sizes (“n_eff”) and split Rhats (“Rhat”) from the fit that used to assess the convergence.

Table S11. Leave-one-out cross validation of the three prior distirbutions using the “loo” package. The expected log posterior densityy (“elpd_loo”) and corresponding standard error (“se_elpd_loo”), and the difference between each prior and the best prior (“elpd_diff”) and the standard error of the difference (“se_diff”). The values of “p_loo” and “looic” and corresponding SE are also included for reference.

Table S12. Summary statistics of lead variants identified in the conditional analyses from sex-combined GWAS and their posterior probabilities within each sex-biased component and their component assignments, as well as sex difference z-score (“SEX_DIFF_Z”) of each variant. “ALLELE1” was used as the effect allele and “ALLELE0” the reference allele in GWAS. The “INFO” contains imputation quality score of variants. Suffixes are used to indicate if the estimation was performed in the sex-combined (“_COMB”), female (“_FEMALE”) or male (“_MALE”) population. “A1FREQ” contains the allele frequency of the effect allele in the sample and “MAF” the minor allele frequency in the sample. “BETA”, “SE”, “CHISQ_BOLT_LMM_INF”, “P_BOLT_LMM_INF” are the estimated effect size, corresponding SE, chi-square statistics, and *P*-values from BOLT-LMM infinitesimal mixed model, respectively. We report the posterior probabilities of each lead SNP in the null effect (“p[0]”), the female-biased effect(“p[1]”), equal effect (“p[2]”) and male-biased effect (“p[3]”) and the assigned component (“COMPONENT”).

Table S13. Summary statistics estimated in FinnGen release 10 of height-associated lead variants identified in UKB sex-combined GWAS and their posterior probabilities within each sex-biased component and their component assignments, as well as sex difference z-score (“SEX_DIFF_Z”) of each variant. “ALLELE1” was used as the effect allele and “ALLELE0” the reference allele in GWAS. Suffixes are used to indicate if the estimation were performed in female (“_FEMALE”) or male (“_MALE”) population. “A1FREQ” contains allele frequency of the effect allele in the sample. “BETA”, “SE”, “P” are the estimated effect size, corresponding SE and *P*-values, respectively. We report the posterior probabilities of each lead SNP in the null effect (“p[0]”), the female-biased effect(“p[1]”), equal effect (“p[2]”) and male-biased effect (“p[3]”) and the assigned component (“COMPONENT”).

### Supplemental Note

#### Motivation and consequence of alternative coding system in chrX

ChrX can also be coded as the count of observed alleles in males and females[1]. Assuming full XCI, a half of the active allele effect is being estimated in females and the full allelic effect in males. Thus, the observed male effects are expected to be twice the female effects when the active allele effect sizes are equal between the sexes (Table S14). In females, $\beta_X = \beta_A$ corresponds to two-fold larger $a_X$ than $a_A$, while in males, $\beta_X = \beta_A$ corresponds to equal $a_X$ and $a_A$. On the other hand, if there was no XCI and both alleles in females were fully functional, then this approach would be estimating per allele effect also in females. Such an approach has been used for studying escape from XCI [2–4]. However, in general, this may be an unrealistic assumption as genes escape from XCI were rarely fully expressed[5].

A biologically intuitive approach is to count each allele in females as 0.5 copies and in males as one copy as implemented in SNPTEST v2.5[6]. This method automatically takes into account full XCI and is estimating active allele effect in both sexes (Table S14). Thus, when comparing with autosomal effect sizes, a similar magnitude of chrX effect would suggest an equal $a_X$ and $a_A$.

Table S14. Expected relationship of metrics between sexes and between chrX and autosomes under other coding systems.

$p$, the minor allele frequency. $var_m$, $var_f$, genotype variance of chrX in male and female analyses, respectively. $\beta_m$, $\beta_f$, male and female effects in chrX, respectively. $var(\beta_m X_m)$, $var(\beta_f X_f)$, phenotypic variance explained by a X-linked variants in males and females, respectively. $h_m^2$, $h_f^2$, chrX heritabilities in females and males, respectively. $a_X$, $a_A$, active allele effects in chrX and autosomes, respectively. $\beta_X$, $\beta_A$, chrX and autosomal effects, respectively. $var(\beta_X X_X)$, $var(\beta_A X_A)$, phenotypic variance explained by chrX and autosomal variants, respectively. $h_X^2$, $h_A^2$, chrX and autosomal heritabilities, respectively. $\phi$, the ratio of the counts of variants contributing to heritability in chrX and in autosomes. Here, $h^2$ is defined as $var(\beta X)/var(Y)$, where $var(Y)$ is the total variance of the trait.

| Genotype coding in chrX non-PAR |
|---|

<table>
<tr><td colspan="4">Male: {0,1}<br>Female: {0,1,2}</td><td colspan="2">Male: {0,1}<br>Female: {0,0.5,1}</td></tr>
<tr><td colspan="6">Genotype variance of chrX variants ($var(X)$)</td></tr>
<tr><td colspan="4">$var_m = p(1-p)$<br>$var_f = 2p(1-p)$</td><td colspan="2">$var_m = p(1-p)$<br>$var_f = 0.5p(1-p)$</td></tr>
<tr><td colspan="6">Assuming full XCI and same active allele effect between sexes in the chrX ($a_f = a_m$):</td></tr>
<tr><td colspan="4">$\beta_{m,X}$ – active allele effect size<br>$\beta_{f,X}$ – half of active allele effect size<br>$\beta_m = 2\beta_f = a_X$<br>$var(\beta_m X_m)/var(\beta_f X_f) = 2$<br>$h_m^2/h_f^2 = 2$</td><td colspan="2">$\beta_{m,X}$, $\beta_{f,X}$ – active allele effect size<br>$\beta_m = \beta_f = a_X$<br>$var(\beta_m X_m)/var(\beta_f X_f) = 2$<br>$h_m^2/h_f^2 = 2$</td></tr>
<tr><td colspan="2"></td><td>X = AA</td><td>X = A</td><td>X = AA</td><td>X = A</td></tr>
<tr><td colspan="2">$a_X/a_A$</td><td>2</td><td>1</td><td>2</td><td>1</td></tr>
<tr><td rowspan="2">$\beta_X/\beta_A$</td><td>male</td><td>2</td><td>1</td><td>2</td><td>1</td></tr>
<tr><td>female</td><td>1</td><td>0.5</td><td>2</td><td>1</td></tr>
<tr><td rowspan="2">$var(\beta_X X_X)$<br>$/var(\beta_A X_A)$</td><td>male</td><td>2</td><td>0.5</td><td>2</td><td>0.5</td></tr>
<tr><td>female</td><td>1</td><td>0.25</td><td>1</td><td>0.25</td></tr>
<tr><td rowspan="2">$h_X^2/h_A^2$</td><td>male</td><td>$2\phi$</td><td>$\phi/2$</td><td>$2\phi$</td><td>$\phi/2$</td></tr>
<tr><td>female</td><td>$\phi$</td><td>$\phi/4$</td><td>$\phi$</td><td>$\phi/4$</td></tr>
</table>

#### Validation of GENESIS

We validated the performance of GENESIS by comparing the estimated $h_A^2$ with the corresponding estimate from LDSC[7], a tool applicable to autosomal data only. Sex-specific estimates from the two methods were well aligned (Pearson $r$ = 0.8 in both sexes) with most traits (36/48 in females and 34/48 in males) showing no detectable differences between the methods (*P*-value ≥ 0.05, two samples *Z* test; Figure S15, Table S2).

#### Number of genome-wide significant variants

We observed a clear sex difference in the numbers of significantly associated X-linked loci. At the genome-wide significance threshold of $5 \times 10^{-8}$, we detected a

2.3-fold (SE = 0.19) the number of associated LD independent variants in males (range from 0 to 25 per trait, median 3) compared to females (range from 0 to 13 per trait, median 1) (Figure S4A, Table S5). These observations did not seem to be driven by sex-specific genetic architecture in chrX as male and female effect estimates of these variants identified from sex-specific analyses were similarly highly correlated in chrX as in the autosomes (Figure S16). The differences in the number of detected associations can rather be explained by greater statistical power in males than in females to detect genetic associations in chrX, a difference arising from the unique biology of chrX. In comparison, in the autosomes, we observed slightly more independently associated genome-wide significant loci in females than in males (1.2-fold (SE = 0.03), Figure S4B), a finding at least partly attributable to the larger sample size of the female subsets for all traits except for direct bilirubin and testosterone (on average 7.62% more females than males; Table S1).

#### The effect of allele frequency and functional consequences on active allele effects

As variants with low frequency and severe functional consequences tend to have large effects, we assessed if the differences in the effect size estimates were explained by differences in MAFs and functional consequences between autosomes and chrX. We first examined how chrX differs from autosomes in general using ~1.1 million independent common variants from the GENESIS 1000 Genomes European reference panel. We observed only minor differences in the allele frequency distribution between chrX and autosomes — variants in chrX have overall a slightly higher MAF (median 0.253 versus 0.251; *P*-value < 0.001, *t*-test), and also a slightly larger proportion of less common variants (0.01 < MAF < 0.05) (1.7% versus 1.4%; *P*-value < $5 \times 10^{-6}$, $\chi^2$ test; Figures S9A and S9B) relative to autosomes (here we note that more than 75% of low frequency variants were with MAF ≥ 0.04). Further testing the relationship of MAF and $a$ at the identified sex-specific trait-associated variants, we observed significantly larger $a_X$ compared to $a_A$ across all the MAF bins (Figures S9C and S9D), suggesting limited effect of MAF on the difference of $a$ between chrX and autosomes.

With regards to variant functional consequences, we observed differences in the active allele effect distributions between chrX and the autosomes. In general, the common variants in chrX were depleted in regulatory and coding regions (predicted by Variant Effect Predictor[8]; Figure S10) and showed enrichment for less pathogenic consequences (estimated by the CADD score[9]; Figure S11A) relative to autosomes. This reduced density of functional variants in chrX aligns with the stronger selection pressure on chrX arising from the hemizygosity of chrX in males[10,11]. Across the trait-associated variants, we again observed larger $a_X$ compared to $a_A$ but this difference was driven by variants with regulatory and other less severe functional impacts whereas no difference between $a_X$ and $a_A$ was observed at the coding region variants (6 (10%) and 226 (4.3%) of the chrX and autosomal lead variants, respectively) (Figure S10B). The same phenomenon was observed when grouping variants based on their pathogenicity. For female lead variants with the greatest pathogenicity (3 (5.0%) and 139 (2.6%) in chrX and autosomes, respectively), we observed no difference between $a_X$ and $a_A$ (median 0.056 versus 0.051, *P*-value = 0.48, Wilcoxon rank-sum test) (Figure S11B); however, for variants predicted as less pathogenic (scaled CADD score ≤ 20), we observed a significantly larger $a_X$ compared to $a_A$ (median 0.048 versus 0.026, *P*-value = $7.09 \times 10^{-18}$, Wilcoxon rank-sum test).

Across all comparisons, the observed patterns were not affected by pleiotropy (Figures. S9E, S9F, S10C and S11C).

#### Dosage compensation ratio analysis

Following the work by [2], we estimated chrX and autosomal DC ratio (DCR) using directly the $\widehat{h^2_{X,m}}$ and $\widehat{h^2_{X,f}}$, $\widehat{h^2_{a,m}}$ and $\widehat{h^2_{a,f}}$ estimated with GENESIS for 34 traits with non-zero $h^2_X$ in both sexes. The DCR and its corresponding standard error were estimated as[2]:

$$DCR = \frac{\widehat{h^2_m}}{\widehat{h^2_f}}$$

$$SE(DCR) = \frac{\widehat{h^2_m}}{\widehat{h^2_f}} \sqrt{\left( \frac{SE^2\left(\widehat{h^2_m}\right)}{\widehat{h^2_m}^2} + \frac{SE^2\left(\widehat{h^2_f}\right)}{\widehat{h^2_f}^2} \right)}$$

where $\widehat{h^2_m}$ and $\widehat{h^2_f}$ are $h^2$ estimates from GENESIS in males and females and $SE\left(\widehat{h^2_m}\right)$ and $SE\left(\widehat{h^2_f}\right)$ are corresponding standard errors. ChrX DCRs ($DCR_X$) were estimated with chrX estimates $\widehat{h^2_{X,m}}$ and $\widehat{h^2_{X,f}}$ with corresponding standard errors and autosomal DCRs ($DCR_A$) with autosomal estimates $\widehat{h^2_{A,m}}$ and $\widehat{h^2_{A,f}}$ with standard errors.

We compared our estimated DCR with DCR estimated using summary statistics as[2]. Our DCR estimates were correlated with that estimated with summary statistics (Pearson $r$ = 0.93 and 0.40 for autosomes and chrX, respectively; Figure S17). DCR estimated with summary statistics had much smaller standard errors and we observed discordant DCR estimates for urate, sex-hormone binding globin, and waist-to-hip ratio (WHR) in autosomes and testosterone in chrX, which may be due to the DCR estimated with summary statistics being less robust to regional sex difference given the lack of consideration of LD.

As sex difference in $h^2$ were observed for autosomes (first section of results), we adjusted the chrX DCR estimates with the corresponding autosomal DCRs to account for sources of sex differences in $h^2$ that are unrelated to chrX specific biology, assuming the effect of e.g. environmental sex biases reflected similarly in autosomes and chrX $h^2$ (Table S2):

$$DCR_{X,adjusted} = \frac{DCR_X}{DCR_A}$$

$$SE\left(DCR_{X,adjusted}\right) = \frac{DCR_X}{DCR_A}\sqrt{\left(\frac{SE^2(DCR_X)}{{DCR_X}^2} + \frac{SE^2(DCR_A)}{{DCR_A}^2}\right)}$$

where $DCR_X$ and $DCR_A$ are estimated DCR in chrX and autosomes, respectively and $SE(DCR_X)$ and $SE(DCR_A)$ are corresponding standard errors in chrX and autosomes, respectively.

For most traits, the adjustment did not introduce overwhelming changes (mean unadjusted 2.46 versus adjusted 2.42; Figure S18) as most autosomal DCR estimates were close to one. The most pronounced change was observed for testosterone (unadjusted 4.67 (SE = 1.92) versus adjusted 2.86 (SE = 1.24)).

We used DCR to test the three XCI scenarios – full XCI (F-XCI), escape XCI (E-XCI) and no XCI (N-XCI). Across the traits, we observed a mean adjusted DCR of 2.40 (SD = 1.15) suggesting, in general, concordance with F-XCI and E-XCI rather than N-XCI, as expected given the existing evidence for XCI[2]. At the individual trait level, while we observed the adjusted DCRs of DBP (0.71 (SE = 0.35) and SBP (0.61 (SE = 0.23)) aligned with the expected value under N-XCI (DCR=0.5), echoing our previous results, the DCR metric did not distinguish between F-XCI (DCR=2) and E-XCI (DCR=1.75) for any of the traits (Figure S19). We observed DCRs greater than 2 for albumin, creatinine, and body fat and mass related traits (body fat mass, weight, impedance of body and left leg), of which correlated traits, body fat percentage and basal metabolic rate, have been reported with DCR greater than 2 due to substantial sex difference in SNP effects in two regions near the *FAM9A/FAM9B* genes and near the *AR* gene[2], that are thought to reflect fat-reducing effects of androgen in males. We additionally calculated DCRs using FinnGen data for height, BMI and weight. While we observed similar DCRs for height as in the UKB, the DCR estimates for weight and BMI differed (Figure S19; Tables S2).

#### Sex-biased effect analysis

##### *Four-component sex bias mixture model*

Demonstrating the consistency of our model, we observed the point estimates of the null effect proportion negatively correlated with $h^2$ estimates in both chrX (Spearman $r$ = -0.40 in females and -0.30 in males) and autosomes (Spearman $r$ = -0.60 in females and -0.63 in males) across traits with nonzero $h_X^2$ in both sexes (Figure S20).

Three different prior distributions of $\sigma^2$ were tested: Inverse-Gamma(1,1), Inverse-Gamma(0.001,0.001) and Uniform(0,1). We compared the estimated parameters with these priors using summary statistics of height for chrX variants. We calculated the expected log-predictive density with leave-one-out cross-validation (ELPD-LOO) for each prior with “loo” R package and compared ELPD-LOO across different priors (Figure S21). The comparison indicated differences between the priors for $\sigma^2$, and ELPD-LOO was the highest for Uniform(0,1) prior. Therefore, we chose to use Uniform(0,1) as the prior for $\sigma^2$ in our analyses.

*Male-biased effects in chrX in testosterone genetics*

Testosterone has been shown in previous research[12–14] to exhibit sex-specific genetic architecture in the autosomes. We observed, as expected, systematically larger effects in males across the genome, however, compared to autosomes, the chrX was more enriched with male-biased variants (scaled proportion 95.3% (95% CI: 85.5 – 99.7%) versus 78.7% (95% CI: 55.9 – 94.6%); Table S9)), an observation consistent with a previous study focusing on sex-specific effects[12]. Such pattern supports the predicted enrichment of variants in chrX that affect traits towards the male optimum[15]. For example, among the 12 lead variants for testosterone in chrX that show male-biased effects, six loci (rs12015400, X:65779624_GTT_G, rs189261721, rs146415516, rs140812443, rs7052964) are close to genes involved in the androgen receptor pathway (*AR* (androgen receptor), *EDA2R* (ectodysplasin A2 Receptor), *KLF8* (KLF transcription factor 8)[16]) and one (rs112265145) close to *FAM9A/FAM9B* region related to spermatogenesis in adults[17] (Table S12). Three regions associated with testosterone showed pleiotropic male-specific effects (Table S12), two of which have been identified as sex-heterogeneous regions[2]: in the *FAM9A/FAM9B* region led by rs112265145 in testosterone association, loci associated with impendence of body, phosphate, heel bone mineral density, insulin-like growth factor 1, total bilirubin, and creatinine display male-specific effects; within the well-known androgen associated locus[2,18,19], *EDA2R*/*AR* region in Xq12, we observed loci associated apolipoprotein A, apolipoprotein B (APOB), high-density lipoprotein, WHR, vitamin D, body fat mass, triglyceride, and creatinine all show considerable larger effects in males except for APOB associated rs35176586 showing slightly larger effects in females; in the *RGAG1*/*CHRDL1* region led by rs881090 in testosterone association in Xq23, a known lipid-associated region[20], we observed lead variants associated APOB, cholesterol, low-density lipoprotein, albumin, sex hormone-binding globulin, aspartate aminotransferase and calcium all display male-biased effects except for X:109833687_GGT_G association with calcium show a female-biased effect.

##### *Lack of female-biased effects in chrX in WHR genetics*

Out of 8 lead associated with WHR in chrX, we identified only two lead variants with female-biased effects – rs113303918 in the intron of *FHL1* (four and a half LIM domains 1) and rs113303918, a missense variant in *SRPX* (sushi repeat containing protein X-Linked) (Table S12), consistent with previous findings in UKB[2]. rs4419961 in the *EDA2R* /*AR* region was identified having larger effects in males on WHR (Table S12).

#### Replicability of sex-biased effects

To understand the poor replicability of the female-biased variants across biobanks, we asked if the sex-specific effects differed between the biobanks. For the sex-combined lead variants for height, we observed strong correlations between UKB and FinnGen in both female (chrX: Pearson $r$ = 0.88, sign test for sign concordance *P*-value = $3.18 \times 10^{-13}$); autosomes: Pearson $r$ = 0.93, sign test *P*-value < $10^{-15}$) (Figures S13C and 14C) and male effects (chrX: Pearson $r$ = 0.92, sign test *P*-value = $3.18 \times 10^{-13}$; autosomes: Pearson $r$ = 0.93, sign test *P*-value < $10^{-15}$) (Figures S13D and S14D), confirming the genetic effects on height in both autosomes and chrX are highly reproducible across data sets. Following these observations, we asked if the sex differences in effect estimates, measured as the sex difference z-scores, correlate between the biobanks. We found these z-scores weakly correlated between the two biobanks, with a small enrichment in directionally concordant effects in the autosomes (chrX: Pearson $r$ = 0.32, sign test for sign concordance *P*-value = 0.78; autosomes: Pearson $r$ = 0.13, sign test *P*-value = 0.004) (Figures S13B and S14B). Overall, we found little evidence of consistency in sex differences in the effect sizes for human height between the two biobanks.
